# Mapping Structural Constraints and Adaptive Potential in a Capsule-Degrading Phage Tailspike Protein

**DOI:** 10.1101/2025.10.06.680778

**Authors:** Sarah Evert, Phill Huss, Dinesh Kumar Kuppa Baskaran, Karthik Anantharaman, Srivatsan Raman

## Abstract

Bacteriophage tailspike proteins (TSPs) degrade bacterial capsules to enable infection, yet the molecular determinants of their function and host range remain unclear. We applied deep mutational scanning (DMS) to the endosialidase TSP of *Escherichia coli* K1 phage K1F, generating 22,365 single-amino-acid variants using an enhanced ORACLE phage engineering platform. Functional scores revealed the TSP is structurally fragile yet harbors pockets of adaptive flexibility. Mutations within the β-propeller active site uncovered residues accommodating longer sialic acid chains than captured by structural studies, while the β-helix stalk emerged as an adaptive “tuning knob” modulating processivity and specificity. Comparative selections across K1 strains identified discrimination hotspots in β-barrel loops and distal residues outside canonical binding sites, implicating capsule modifications and O-antigen presence as key modulators of host range. By resolving how specific mutations modulate function and host range, this study offers a roadmap for designing phages that overcome capsule-based defenses in pathogenic bacteria.

## Introduction

Bacteriophages (phages) infect bacteria with remarkable host specificity and play central roles in shaping microbial ecosystems. This specificity has made them attractive candidates for targeting antibiotic-resistant pathogens that are otherwise difficult to treat^1–4^. The host specificity of phages is primarily mediated by receptor binding proteins (RBPs) located at the phage tail fiber or tail spike, which mediate host recognition and adsorption to diverse ligands on the bacterial surface. These proteins are remarkably adaptable, enabling phages to target a wide array of host species despite rapidly evolving surface architectures^5–8^. The best-studied RBPs bind exposed outer membrane components such as lipopolysaccharides (LPS) or proteins^5, 9–11^. However, numerous clinically important bacteria are encased in dense, strain-specific capsules composed of capsular polysaccharides (CPS), highly variable carbohydrate structures that form a protective barrier around the cell^12–15^. These capsules can sterically shield underlying receptors, rendering conventional RBPs ineffective and significantly restricting phage infectivity^16–20^. Phages have overcome this barrier by deploying a specialized class of enzymatic RBPs, known as tailspike proteins (TSPs), which unlike other RBPs, are architecturally and functionally more complex^16, 21, 22^.

These unique TSPs are catalytically active molecular machines that both bind specific glycan motifs within the capsule and degrade them, effectively “drilling” through the capsule to access the bacterial surface^16^. This dual functionality, high affinity molecular recognition coupled with targeted degradation, distinguishes TSPs from other RBPs and makes them critical determinants of infectivity against encapsulated bacteria. Capsular sugar chemistries are extremely diverse, even within one bacterial strain, resulting in TSPs with diverse substrate specificities and catalytic strategies^12, 23–25^. Each TSP is exquisitely tailored to its hosts capsule architecture and has co-evolved with their bacterial hosts to accommodate this broad array of capsule chemistries, similar to what is observed for other classes of RBPs^16, 20, 25–27^. While conventional RBPs have been extensively studied, capsule-degrading TSPs remain vastly underexplored, even though they are often the decisive factor in whether a phage can breach the capsule barrier and infect clinically important pathogens^28–31^.

To date, much of what is known about TSPs has been derived from structural and biochemical studies using purified proteins and substrates often consisting of single saccharide units. These efforts have yielded critical insights into TSP architecture, glycan specificity, and catalytic mechanisms, but they leave important gaps. Specifically: (1) they rely on purified proteins and simple mono-or short oligosaccharides, which do not reflect how complex capsular sugars are presented on the bacterial surface; (2) they overlook the complexity of phage infection, which occurs in heterogeneous environments where capsule density, glycan presentation, and accessory surface structures can modulate activity; and (3) they do not capture the multifunctional nature of TSPs, in which recognition and catalysis are integrated within a single protein, nor do they interrogate how individual mutations reshape activity and host range^20, 32–38^

For conventional RBPs, single point mutations have been seen to profoundly alter their function. For example, a single mutation in the phage Φ21 RBP enables recognition of a new outer membrane protein, while single point mutations to the T7 RBP can increase infectivity on diverse hosts^39, 40^. These findings suggest that TSPs, which must both bind and enzymatically degrade capsule glycans, may be even more sensitive to mutational perturbations. Yet no systematic maps exist of how TSP mutations impact glycan recognition, catalytic efficiency, or host range in the native infection context. Given their enzymatic complexity and the extended capsule interface they engage, only a comprehensive approach such as deep mutational scanning (DMS) can reveal the full spectrum of sequence–function relationships that govern TSP activity and determine phage infectivity against encapsulated bacteria.

Here we apply DMS of the K1F phage TSP to uncover molecular determinants driving activity and host range^41^. K1F is a model *Escherichia coli* K1 phage whose endosialidase TSP specifically recognizes and cleaves the α-2,8-linked polysialic acid chains of the K1 capsule, allowing the phage to breach this barrier^42–44^. Using an enhanced version of the ORACLE phage engineering platform, tailored to accommodate large and functionally complex genes such as TSPs, we generated a comprehensive library of 22,365 single amino acid substitutions spanning the full 3.2 kb TSP gene^40^. We then systematically dissected the contribution of each residue to catalysis and host range by quantifying variant function across multiple bacterial hosts. This approach yielded high-resolution functional maps that revealed (1) unexpected gain-of-function mutations both near and far from substrate-binding sites, (2) novel catalytic residues outside the canonical active site, and (3) context-dependent determinants that differ between *in vitro* assays and native phage infection. In addition, we uncovered permissive regions that modulate host discrimination even among strains with identical CPS serotypes and identified functional determinants outside the canonical CPS synthesis locus, revealing that phage–host specificity is shaped by previously unrecognized cell surface features.

Together, these findings define a blueprint for decoding and engineering enzymatic RBPs, establishing a broadly applicable framework for rational phage design against encapsulated bacterial pathogens

## Results

### Construction of a Comprehensive DMS Library of the K1F Tailspike Protein

We successfully constructed an 18,886-member deep mutational scanning (DMS) phage library of the entire K1F tailspike protein, encompassing all 19 possible non-synonymous substitutions and a stop codon at each codon position, using an enhanced version of the high-throughput phage engineering method ORACLE^40^. ORACLE involves four key steps; acceptor phage generation, Cre-mediated recombination from a donor plasmid, enrichment for recombined phages, and library expression prior to selection^40^ (Supp. 1). Originally, ORACLE was demonstrated on a much smaller (∼500 bp), non-enzymatic target gene, where direct sequencing was feasible, donor plasmids were stable, and recombination efficiency was high. Scaling to a gene three times larger and encoding a complex enzymatic protein required two key innovations: (i) implementing a DNA barcoding strategy to track variants after genome insertion, and (ii) stabilizing donor plasmids by relocating recombinase under inducible genomic control, which together improved recombination efficiency by an order of magnitude. These advances enabled construction of a comprehensive DMS library for functional mapping of the K1F TSP in its native phage context, establishing the feasibility of extending ORACLE to larger genes (1A).

To build the TSP DMS library, we first constructed a barcoded plasmid library carrying all variants for phage genome insertion. The gene was divided into 20 segments, with each sub-pool assembled from oligonucleotides containing all relevant mutations and an adjacent random DNA barcode for barcode-variant mapping via separate mapping constructs^45^. The resulting 20-subpools were then combined to generate the final 22,365 member variant library (1B). On average, the library contained ∼200 barcodes per variant, though coverage varied across pools (e.g. pool 16 average 50 barcodes per variant, pool 18 average 146, and pool 1 averaged 362) (Supp 2A-B). Despite this variation, coverage was sufficient to proceed with phage genome insertion. The final barcoded plasmid library represented 95.2% of the full library (21,289 variants of a possible 22,365 member library).

Next, we employed an enhanced version of the previously developed method, ORACLE, to insert library variants into the K1F genome. As outlined above, we made key changes to the recombination step to accommodate large and complex gene insertions, including introducing DNA barcoding to track variants after genome integration and relocating the recombinase under inducible control to stabilize donor plasmids and boost recombination efficiency.

For library integration, we first constructed a K1F acceptor phage in which the K1F TSP gene was replaced with a fixed DNA insert flanked by Cre recombinase sites, following the original ORACLE design^40^. This acceptor phage exhibited no significant plaquing deficiency compared to wild type K1F when the TSP was supplied *in trans* (Supp. 3A). Importantly, introducing Cre recombinase sites directly adjacent to the wild type TSP did not impair phage fitness (Supp. 3B).

Next, we attempted to insert library variants into the phage genome using Cre recombinase. However, plasmids carrying the K1F TSP together with constitutive Cre (original ORACLE method) were unstable, likely due to the metabolic burden of co-expressing two large proteins during recombination. To overcome this, we engineered a “recombination” strain of *E. coli* EV36 with inducible Cre integrated into the genome, which did not affect phage viability (Supp. 3C). By optimizing induction conditions in this strain, we increased recombination efficiency ten-fold, from ∼1 in 10,000 to ∼1 in 1000 phages, reaching levels comparable to the original ORACLE system^40^ (Supp. 3D).

Following recombination, we enriched for recombinant phages using a Cas9/gRNA system targeting the fixed acceptor sequence, as in the original ORACLE workflow. This system successfully eliminated unrecombined phages without off target effects and within ∼90 minutes, the total population was nearly ∼100% recombined phages (1D, Supp. 4A-B). A final passage on the host allowed expression of the integrated TSP variant off the phage genome for the first time. The resulting library retained 18,866 variants (85% of the designed 22,365), with strong correlation (R^2^=0.826) between independent assembly replicates, underscoring the robustness of the approach (Supp. 4C-D).

The observed loss of variants during ORACLE could reflect (i) insufficient recombination efficiency, (ii) unintended selection during the workflow, or (iii) negative selection against structurally disruptive mutations. In a test library of 2,268 members, the same variants consistently dropped out across ORACLE replicates, with reproducible losses at specific positions, indicating the bias was not due to recombination efficiency (Supp. 5A). Constructing the same library in *E. coli* 10G, which K1F cannot infect, produced similar dropout patterns, arguing against host-driven selection (Supp. 5B). Together, these results point to intrinsic structural constraints, with certain mutations so disruptive that they cannot be tolerated even in the presence of the wild type protein.

Overall, we demonstrate that the enhanced ORACLE can build phage variant libraries for much larger and functionally complex proteins than previously reported. By incorporating DNA barcoding to track variants, minimizing plasmid instability, and fine-tuning recombination, we overcame key challenges associated with scaling to the 3.2kb K1F TSP. These innovations enabled us to recover a comprehensive DMS library encompassing >18,000 unique variants, establishing the largest functional library of a phage TSP. More broadly, this improved workflow provides a generalizable framework for building variant libraries of other large phage TSPs.

### Structural Fragility and Hidden Adaptive Potential in the K1F TSP

To assess how the K1F TSP tolerates sequence variation and to identify regions of functional constraint and evolvability, we evaluated our DMS library on the model *E. coli* K1 strain, EV36. Functional scores (Fn) were calculated from the relative abundance of each variant before and after selection, normalized to wild type, following approximately two replication cycles (see methods). Fn values obtained by barcode sequencing correlated with direct sequencing of a ∼250 bp sub-pool, with both approaches reporting similar variant activity trends (Supp. 6A). Biological replicate functional scores were also highly correlated, providing confidence in reproducibility (Supp. 6B). As expected, catalytic residues previously reported in literature (H350, R549, and E581) were functionally critical on this host and tolerated very few substitutions while stop codons and start codon mutations were generally not tolerated^32, 46^ (1E, Supp. 6C). Together these results gave us confidence in the functional scores reflected the underlying biology of phage-host interactions.

Next, we asked how this high-resolution dataset could reveal region-specific responses to amino acid substitutions and how these patterns reflect the protein’s architecture and biological role (Supp. 6C). The endosialidase TSP is essential for K1F infection, where it binds and cleaves the long α-2,8 linked polysialic acid chains of the K1 capsule^44, 47^. It is a homotrimeric protein composed of three domains: an N-terminal binding domain that attaches the protein to the phage capsid, a central catalytic domain that binds and cleaves the capsule, a C-terminal chaperone domain (CTD) required for proper folding and is cleaved off upon maturation. Structural studies of the catalytic domain, solved both in apo and substrate-bound states, have revealed three polysialic acid binding sites^32, 43, 48^. Based on these studies, the catalytic domain can be subdivided into the binding sites within the β-propeller active site, the β-barrel, and in the β-helix stalk (Fig. 2A). Separately, the structure of the CTD in complex with the β-helix stalk has been solved further revealing regions important for proper folding of the TSP (2A)^49^. We leveraged this structural framework to interpret patterns of mutational tolerance.

**Figure 1.**
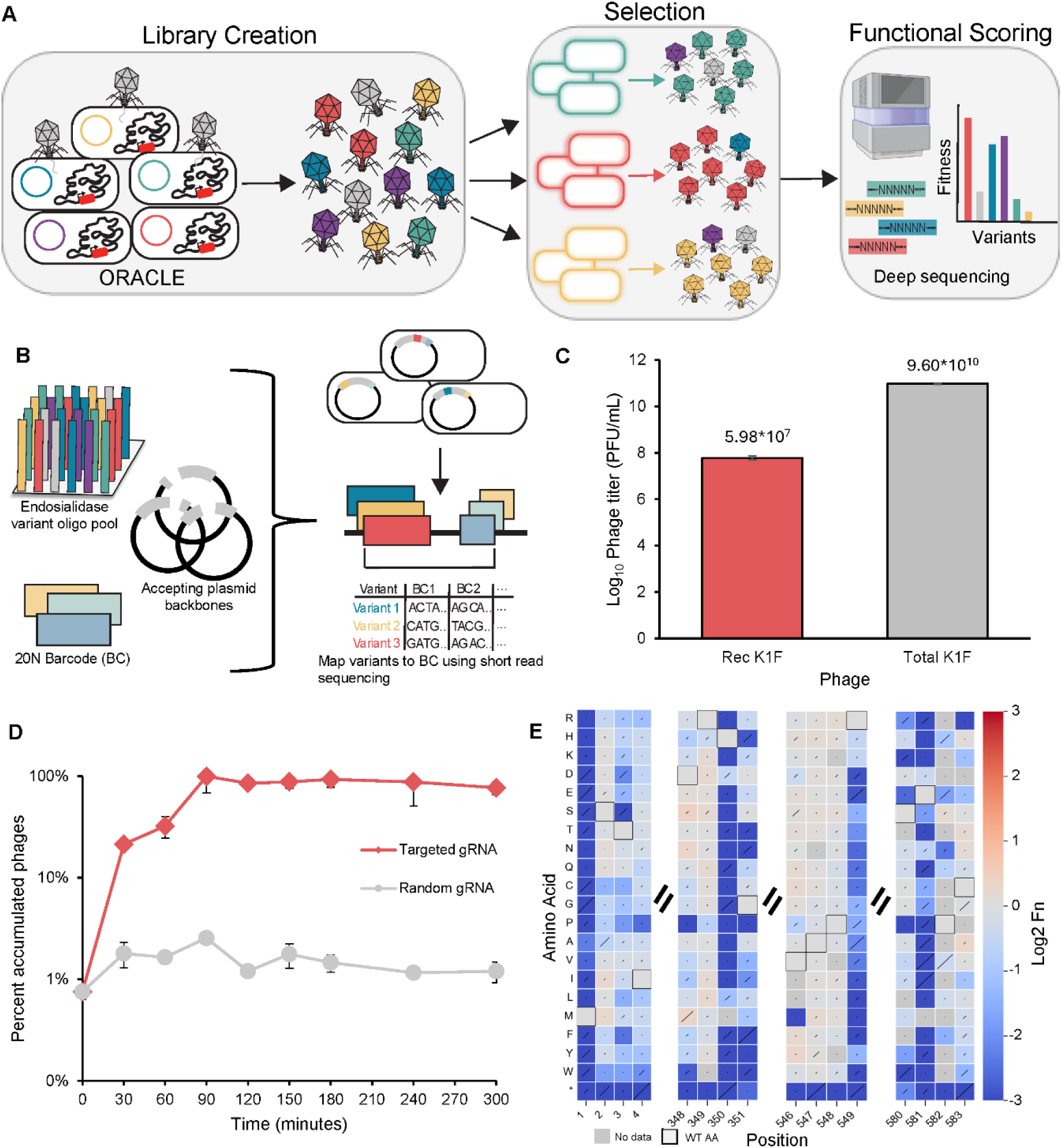
Workflow to build a phage K1F deep mutational scanning (DMS) library using the phage engineering method ORACLE. (**A**) Schematic of the library creation, selection, and sequencing workflow for determining functional hot spots across the K1F tailspike protein. (**B**) Overview of the DMS plasmid barcode mapping process. (**C**) Titers of recombined K1F phage versus total K1F phage following the Optimized Recombination step of ORACLE during full library assembly. Rec K1F is the recombined K1F phage and Total K1F represents both the acceptor phage and recombined phage K1F population. (**D**) Percent of recombined phages in the total population over time during the Accumulation step of ORACLE using a gRNA targeting the randomized DNA sequence of the acceptor phage and random gRNA. (**E**) Heatmap chunks of positions expected to be depleted following selection on *E. coli* K1 strain EV36 for barcode mapping validation. H350, R549, and E581 are previously reported catalytic residues and very few variants are tolerated following selection. Limit of detection is-3.

**Figure 2.**
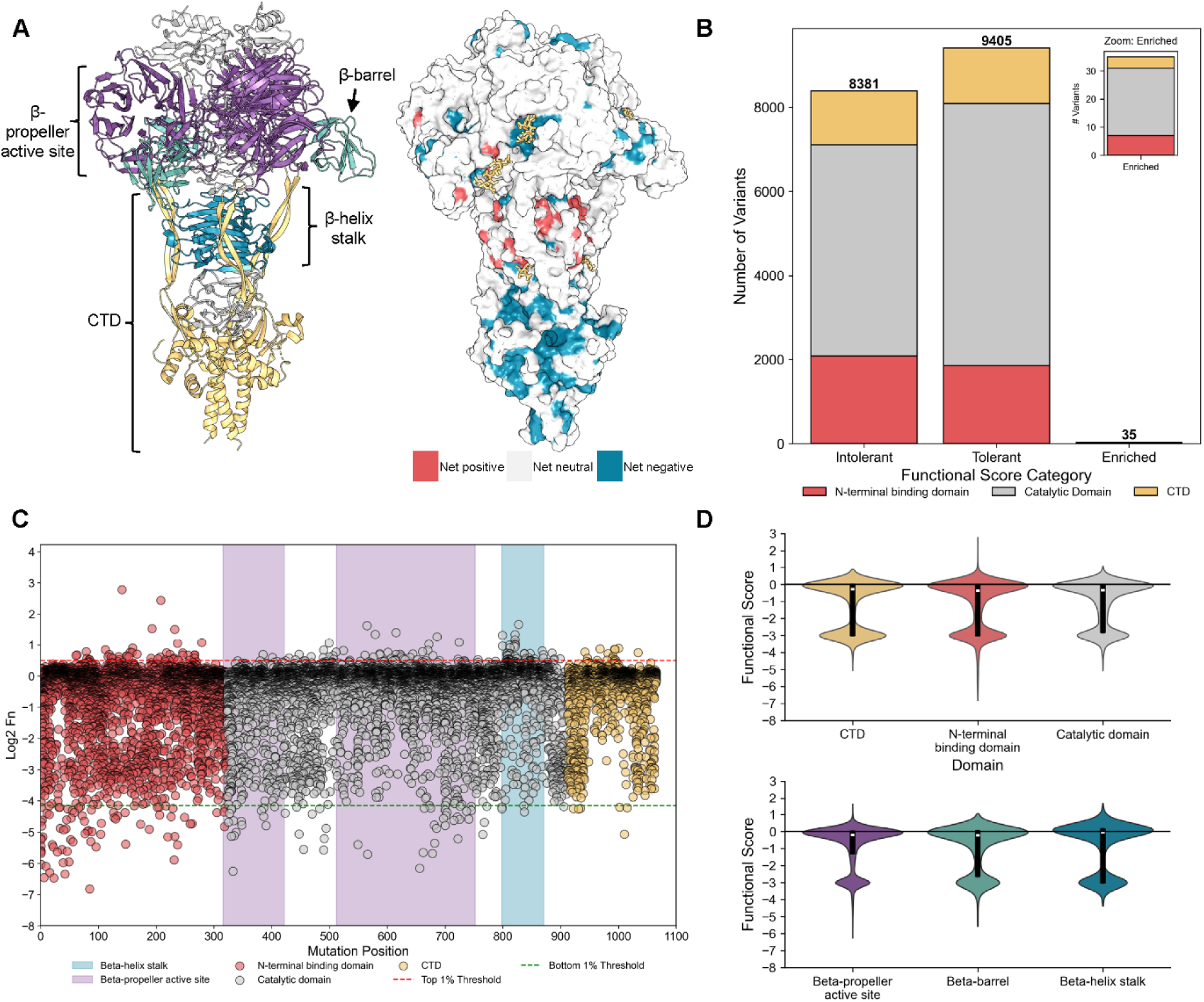
**Distribution of K1F tailspike protein (TSP) variants following selection on *E. coli* EV36**. (**A**) Cartoon representation (left) and surface representation colored by net functional scores at each position (right) of the K1F TSP. Cartoon structure is colored by different domains/sites of the protein (PDB 3gvk and 3gw6 merged). (**B**) Distribution of intolerant (Fn <-0.3), tolerant (Fn ≤ 0.8), and enriched (Fn > 0.8) variants following section on *E. coli* EV36 colored by domain. Includes variants dropped out following selection with the limit detection of-3. (**C**) Scatterplot of the functional scores of all variants at each position colored by domain and shaded by site across the K1F TSP following selection on *E. coli* EV36. Top 1% and bottom 1% are the top 1% and bottom 1% performing variants in the entire library. Dropouts are excluded. (**D**) Violin plots of functional scores for variants in different sites across the K1F TSP. C-terminal chaperone domain (CTD).

Most substitutions are deleterious, though isolated gain-of-function variants emerge, particularly in the β-propeller and β-helix stalk. Nearly half of TSP variants (44%) were depleted after selection, with 24% dropping out entirely, highlighting the structural fragility of the protein (Fig. 2B, Supp. 7A). Variants absent after selection but present in the input library were assigned an Fn score of-3, reflecting the limit of detection. Only 35 variants were enriched (Fn > 0.8, 1σ above wild type distribution) suggesting that the wild type sequence is already highly optimized for infection on EV36 (Fig. 2B, Supp. 7B). Domain-level comparisons revealed broadly similar patterns of mutational constraint. All domains were generally intolerant to substitution, although a subset of N-terminal variants showed enhanced fitness, and certain CTD variants showed broader mutational tolerance (Fig. 2C-D, Supp. 7C). Within the catalytic domain, mutational tolerance varied by structural element and binding site. The β-barrel was largely permissive, with few variants strongly depleted, whereas both the β-propeller active site and β-helix stalk displayed more bimodal mutational tolerance, with many deleterious substitution but also several that improved fitness (Fig. 2C-D, Supp. 7C). These trends motivated a closer examination of mutational tolerance within each domain.

Of 5,883 possible variants in the β-propeller active site, 42% (2,502) were intolerant (Fn ≤ –0.3, >1σ below wild type distribution) (Fig. 3A). Intolerant positions were concentrated in the central channel, responsible for sialic acid coordination and cleavage, whereas peripheral loops were broadly tolerant. Key catalytic residues (H350, R549, E581, R596, and R647) previously implicated in sialic acid binding and cleavage were highly intolerant to substitution^32, 46^. Positions deeper in the channel, such as K585 located 19 Å from the substrate, also tolerated few mutations, suggesting that the active site accommodates longer sialic acid polymers than the trimers inferred from structural studies^50–52^. Conversely, K410, reported as essential *in vitro*, tolerated substitutions and was not essential in phage-based screens, underscoring that requirements inferred from biochemical assays may not always hold in the context of infection^32^ (Fig. 3A).

**Figure 3.**
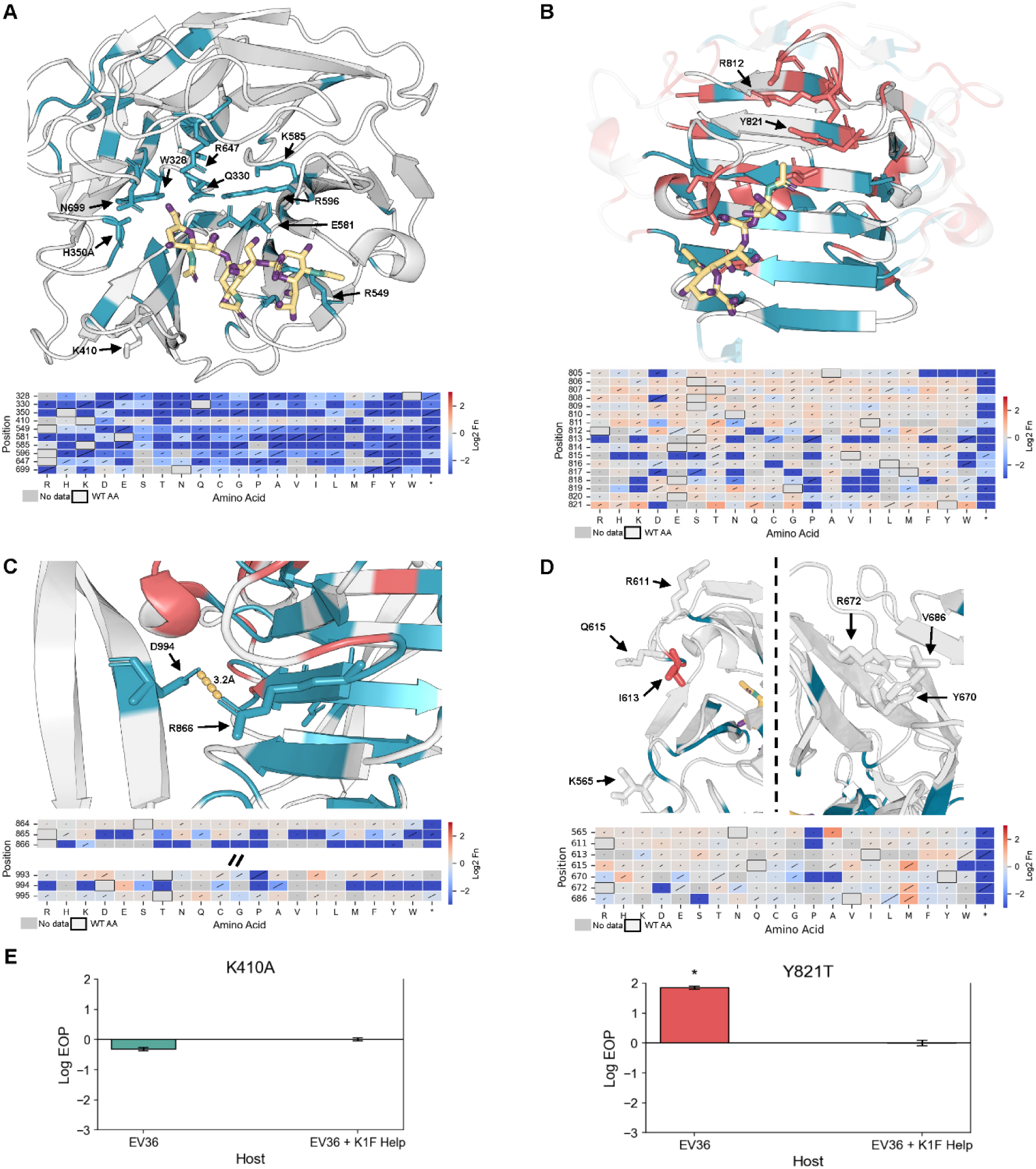
Net impact of all variants at each position following K1F tailspike protein (TSP) library selection on *E. coli* EV36 and resulting discovered functional hotspots. (**A-D**) Cartoon representation of the β-propeller active site, β-helix stalk domain, C-terminal chaperone domain, and peripheral loops of β-propeller active site, respectively. Positions are colored by net normalized functional score (Fn) impact, with negative impact colored blue, positive impact red, and neutral impact gray. Key functional positions are shown in a licorice representation. Corresponding heatmaps of Fn for key positions highlighted in the structure are shown below each structure. Diagonal lines in square represent standard deviation. Hydrogen bonding interactions are represented by gold dashed lines. Polysialic acid is colored in gold, nitrogen green, and oxygen purple. PDB: 3gvk and 3gw6. (**E**) Efficiency of plating (EOP) assay results for two phage K1F clonal variants done in replicate. EV36 + K1F Help is the reference host. *p-value < 0.05.

Building on these observations, we identified previously uncharacterized residues Q330, K585, and N699 in the β-propeller core that exhibited strong loss-of-function phenotypes despite lacking direct substrate contacts in prior structural models (Fig. 3A). N699 can form a hydrogen bond with the substrate suggesting a role in substrate stabilization (Supp. 8A). Q330 and K585 appear to be forming a network of electrostatic and hydrogen-bond interactions with positions D331, H527, S528, and E581, likely stabilizing the catalytic center or coordinating longer sialic acid chains (Supp. 8B). Notably, these residues are also highly conserved across other endosialidases, underscoring their potentially critical role in enzymatic function, despite not being close to substrate binding sites (Supp. 9).

In contrast to the β-propeller, which tolerated many substitutions outside the active site channel, the β-helix stalk showed high mutational plasticity and harbored the greatest density of gain-of-function variants (Fig. 3B, Supp. 7C). Enhanced function was associated with mutations that weakened interactions between the TSP and the sialic acid chain (Supp. 10). Residues normally involved in coordinating the negatively charged sialic acid chains, such as aromatic or positively charged residues, were frequently substituted with hydrophobic or negatively charged residues that improved variant performance. For example, changes from aromatic residue Y821 to smaller or more hydrophobic residues produced variants with enhanced activity, while substitutions of positively charged R812 with hydrophobic or even negatively charged residues also improved variant performance (Fig. 3B, Supp 10). These findings identify the stalk as an adaptive “tuning knob” that modulates substrate processing, providing the direct functional evidence that this region directly shapes phage activity^32, 43^.

The CTD exhibited strong intolerance to substitutions, reflecting its essential role in protein folding and maturation, but tolerated mutations in β-strand extensions that appear structurally peripheral^48, 49, 53^. Notably, only one position, D994, showed tolerance for fewer than 50% of possible mutations, suggesting that it is involved in stabilizing the β-helix stalk domain during folding via hydrogen-bonding interactions with position R866 (Fig. 3C, Supp 6). By contrast, the tolerance of nearly all other β-strand extension positions suggests they may be functionally dispensable for CTD folding. Although the CTD is not well conserved across endosialidase TSPs, both R866 and D994 are conserved, pointing to a critical role for this interaction in protein function (Supp. 9).

We also identified distal residues that exhibited mutational flexibility and signs of evolvability. Several surface-exposed positions extending outward from the TSP in the peripheral loops of the β-propeller active site such as Q615, N565, and V686 tolerated chemically diverse substitutions that improved fitness (Fig. 3D). For example, at Q615, methionine and arginine substitutions were enriched upon selection (Fn = 1.41 and 0.75 respectively), while at N565 and V686, alanine and methionine substitutions were strongly enriched (Fn = 1.63 and 1.02 respectively). Because these residues lie outside the known sialic acid binding pockets, they may engage in transient interactions with other glycans (e.g., O-antigen or LPS) as the TSP drills through the capsule. These mutationally tolerant sites may provide a foothold for adaptive flexibility against diverse host surface chemistries.

Finally, we confirmed that the activity profiles we observed in the pooled DMS screen can be recapitulated through clonal testing of two variants, K410A and Y821T. K410A showed no significant growth defect on EV36 in efficiency of plating (EOP) assays or growth assays, consistent with its dispensability in the native phage context (Fig. 3E, Supp. 11). By contrast, Y821T conferred a significant fitness increase on EV36, supporting the idea that destabilizing stalk interactions can enhance activity (Fig. 3E, Supp. 11). Together, these clonal experiments reinforce the predictive power of our screen.

Overall, our DMS screen reveals that the K1F TSP is structurally fragile yet harbors pockets of adaptive flexibility. Roughly two-thirds of variants were intolerant, underscoring the strong structural constraints across all domains. However, specific substitutions at both binding and distal sites enhanced function, indicating that the TSP is tunable and may engage additional bacterial surface glycans. Notably, residues critical in the native phage context did not always align with those identified in vitro, highlighting the importance of studying TSPs in their physiological environment. Together, these findings show how DMS can uncover both constraints and opportunities for evolvability within a complex enzymatic RBP. DMS of K1F TSP uncovers deep structural fragility but also hidden adaptive potential, highlighting novel targets for phage engineering.

### Cross-Host DMS of K1F TSP Reveals Molecular Basis of Phage Discrimination

The power of a comprehensive mutational library is that it can be deployed across multiple hosts, with each selection experiment producing a high-resolution molecular blueprint of how the phage engages that host. To dissect the determinants of host discrimination, we passaged the K1F DMS library on four additional *E. coli* K1 strains and used unsupervised hierarchical clustering to compare mutational landscapes. Although the α-2,8–linked polysialic acid capsule was thought to be sufficient for K1F infection, the phage must navigate a far more complex environment in the native context, where glycan presentation and accessory surface structures can modulate specificity^47, 54, 55^. Comparing functional landscapes across strains allowed us to pinpoint regions of the TSP that govern both conserved activity and host-specific interactions, providing new insights into the molecular basis of phage discrimination.

The four additional screened *E. coli* K1 bacteria strains were clinically relevant strains from the CDC *Enterobacterales* Carbapenem Breakpoint (BIT) panel, ATCC antibiotic resistance panel, and from clinical urinary tract infection (UTI) isolates^56^.^56^. These strains, UTI50, *E. coli* O45:H10 (ATCC BAA-2649), A204 (SAMN04014854), and A208 (SAMN04014858), can be readily lysed by both wild type K1F and the K1F DMS library (Supp. 12A). These comparative selections provided the opportunity to identify regions of the TSP that vary in their contribution to host specificity.

To pinpoint TSP positions involved in host discrimination, we clustered variants by activity across hosts using correlation-based agglomerative hierarchical clustering (Fig. 4B). We used correlation-based agglomerative hierarchical clustering because of its tolerance of outliers and ability to resolve smaller clusters. Variants with similar activity were grouped into 50 clusters, chosen as the best balance between capturing discrimination trends and avoiding over-fragmentation (Supp. 12C). This partitioning captured major patterns of host discrimination. Importantly, the number of clusters does not affect the underlying relationships between each phage variant and host (Supp. 13A).

**Figure 4.**
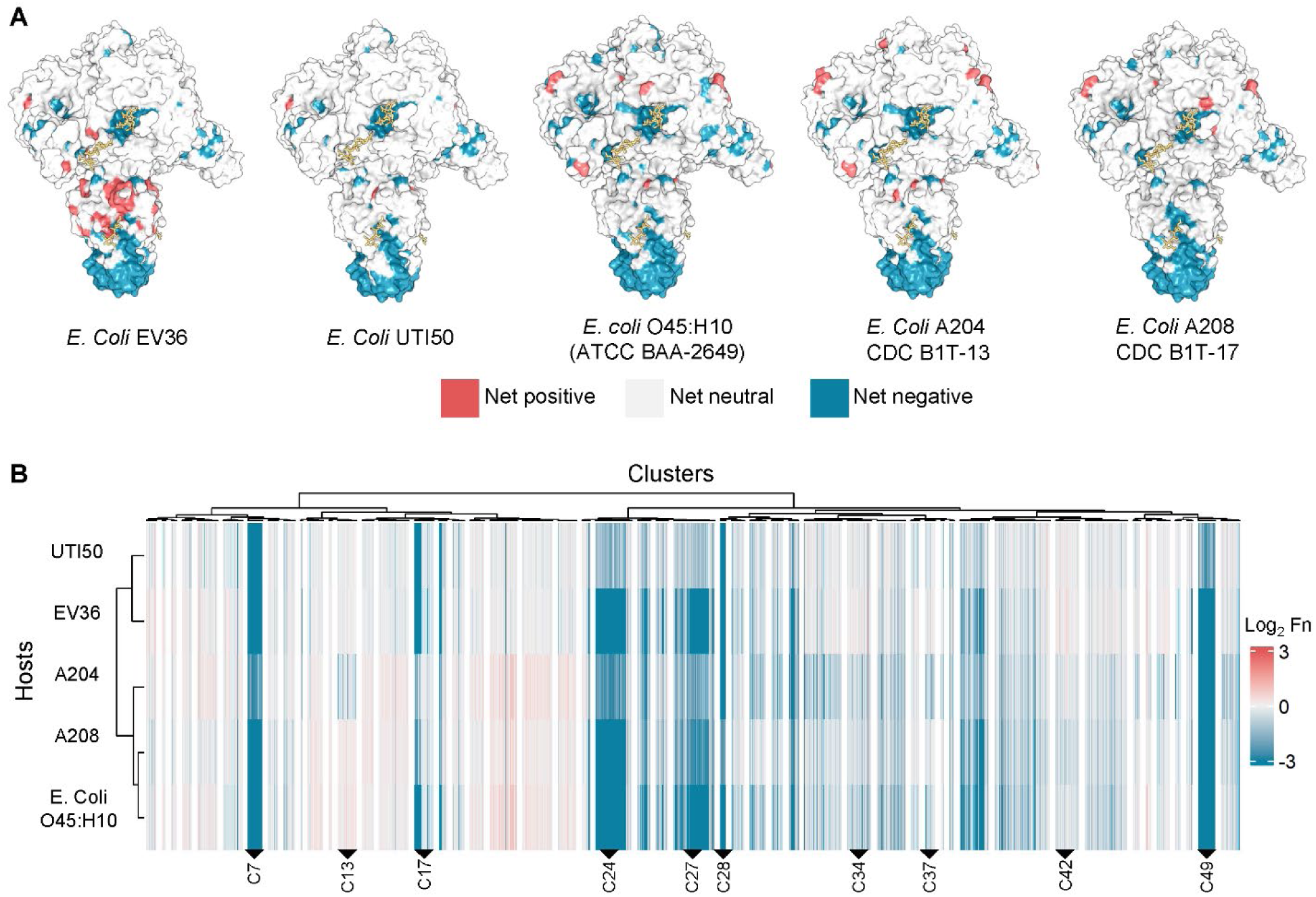
**Selection of the K1F deep mutational scanning (DMS) library across five multidrug bacterial strains and unsupervised hierarchical clustering reveals discriminatory regions**. (**A**) Net impact of each position following selection across 5 bacterial strains mapped onto the K1F tailspike protein (PDB: 3gvk). The polysialic acid chains are color gold in licorice representation. (**B**) Hierarchical clustering heatmap for functional scores following selection of the K1F DMS library on each of the 5 bacterial strains. Key clusters are labeled and there are 50 total clusters represented.

Examination of the hierarchical clustering revealed broadly neutral (similar to wild type across all hosts), disabling (intolerant across all hosts), and host discriminating clusters (Supp. 13B). The latter contained variants that performed better or worse than wild type on at least one host relative to others, pointing to residues with host-specific effects. Strikingly, these host discriminating variants mapped to multiple, distinct regions of the TSP, indicating that host selectivity arises from contributions across the protein. Below, we highlight four examples that illustrate how mutations in different structural contexts shape host selectivity.

Variants in cluster 37 showed fitness deficits specifically on only *E. coli* O45:H10 and A204 (Fig. 5A). These substitutions were predominantly bulky or charged residues replacing smaller or hydrophobic ones (Supp. 14A). The variants with the largest difference in functional scores (top 10%, 20/195 total variants) mapped to positions G730, G742, and N732, all located at flexible loops behind the sialic acid β-barrel binding site (Fig. 5A). Because these mutations lie near a sialic acid binding site, the results suggest that the K1 capsule sialic acid chains of *E. coli* O45:H10 and A204 contain structural modifications, such as O-acetylation, that are sterically hindered when bulkier substitutions are present^57, 58^.

**Figure 5.**
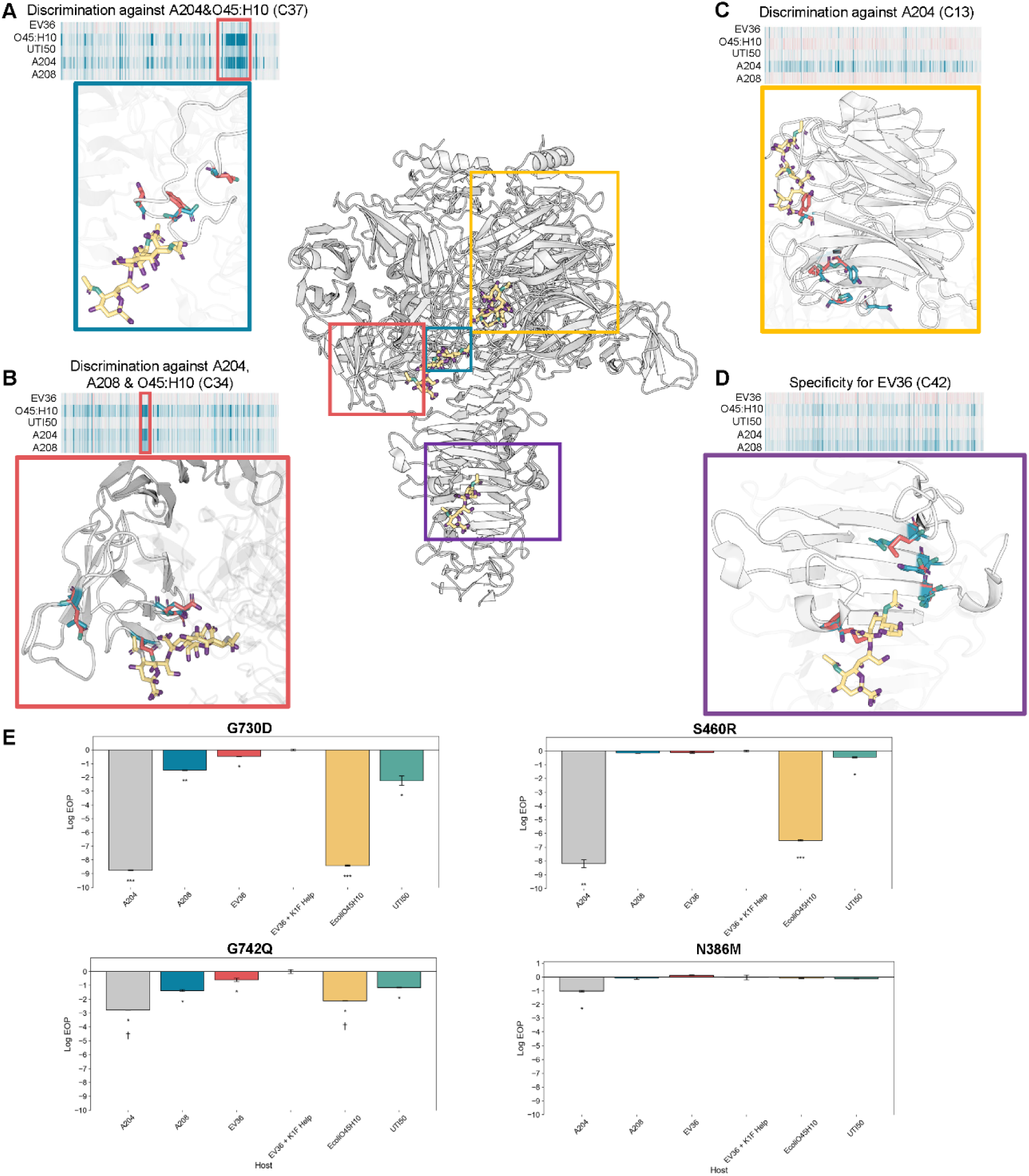
Distinct host discrimination patterns across hosts can be observed following hierarchical clustering. (A) Heatmap for cluster 37. Red box highlights most discriminatory variants. Variants with the greatest discrimination from *E. coli* O45:H10 and A204 versus the rest of the hosts are represented in the structure. The wild-type residue is colored blue, and a discriminatory mutation colored in red. Polysialic acid chain is colored gold, oxygen purple and nitrogen green. (B) Heatmap for cluster 34. Variants with the greatest discrimination on *E. coli* O45:H10 and A204 versus the rest of the hosts are represented in the structure and boxed in red. (C) Heatmap for cluster 13. Variants with the greatest discrimination on A204 versus all other hosts are represented in the structure of the β-propeller. (D) Heatmap for cluster 42. Variants with the greatest discrimination for EV36 are represented in the structure of the β-helix stalk domain. (E) EOP assays for four representative host discriminatory variants across all 5 hosts. EV36 + K1F Help is the reference host. A p-value < 0.05 has *, p-value < 0.01 has **, and a p-value < 0.001 has ***. A cross indicates smaller plaque sizes. PDB: 3gvk.

Cluster 34 variants showed a similar pattern of host-specific depletion, particularly on E. coli O45:H10, A204, and A208 (Fig. 5B). The strongest discriminatory effects arose from substitutions of small or polar residues (e.g., serine or glycine) with negatively charged or hydrophobic residues (e.g., glutamate or valine), including S460R, S468D, and G504V (Supp. 14A). Within the β-barrel binding site, these small or polar amino acids normally accommodate the negatively charged carboxylate groups of the sialic acid and provide flexibility to the binding pocket^59, 60^. Their replacement with bulky or charged side chains led to discrimination on only certain hosts, indicating that differences in the helical conformation of sialic acid chains underlie host-specific effects^61, 62^. A tighter, more compact chain may better tolerate such substitutions, whereas a looser conformation may be sterically hindered, resulting in host-specific discrimination.

Cluster 13 variants were specifically depleted on strain A204 and carried substitutions located away from the sialic acid binding sites (Fig. 5C). These mutations mapped to the periphery of the β-propeller active site (∼18-26Å away) and involved polar to hydrophobic substitutions such as N386M, E373G, and Y387S (Supp. 14A). Because these substitutions lie outside the binding site, they are unlikely to interact directly with the substrate. Instead, they may indirectly effect the active site structure or participate in transient interactions with other bacterial surface sugars. Thus, residues outside canonical binding sites can also shape host discrimination.

Finally, cluster 42, enhanced activity only on EV36 (Fig. 5D). Mutations in this cluster were located at the β-helix stalk, a region we previously found to contain many enriched variants upon selection on EV36. We had proposed that these mutations destabilize interactions with the sialic acid substrate (Supp. 14A). Although destabilization of these interactions is detrimental on some hosts, it appears beneficial on EV36, again suggesting that the β-helix stalk functions as an adaptive “tuning knob” for host specificity.

To confirm these findings, we generated clonal phage variants for key positions from each representative cluster to validate strain-specific effects. In the pooled data, bulky substitutions at G730 and G742 (cluster 37) were deleterious on *E. coli* O45:H10 and A204 but tolerated on the other hosts. EOP validation of clonal variants for G730D and G742Q showed significant growth defect only on *E. coli* O45:H10 and A204, as expected (Fig. 5E, Supp. 14B). The clonal variant S460R (cluster 34) showed significant defects exclusively on A204 and *E. coli* O45:H10, recapitulating the pooled data (Fig. 5E, Supp. 14C). Clonal variant N386M (cluster 13) produced a subtle but a significant growth defect only on A204 similar to the pooled results (Fig. 5E, Supp. 14C) These results corroborate the DMS findings and support the conclusion that these residues contribute to host-specific discrimination.

Together, these comparative selections reveal that host discrimination in the K1F TSP is encoded across multiple structural contexts, from flexible loops adjacent to binding sites to peripheral and stalk regions distant from the catalytic pocket. Rather than a single determinant, specificity emerges from a distributed network of residues whose effects vary depending on host surface chemistry. By using DMS as a comparative tool, we generate molecular blueprints of phage–host interactions that pinpoint residues critical for discrimination and highlight adaptive hotspots. These findings provide a path toward rational engineering of phages with tailored host ranges and illuminate how enzymatic RBPs navigate the complex environments presented by diverse bacterial surfaces.

## Discussion

Phages with depolymerizing tailspike proteins (TSPs) are promising therapeutics against encapsulated and biofilm-forming bacteria, which are notoriously resistant to conventional antibiotics^21, 63–65^. Yet, the molecular determinants that govern how TSPs engage their hosts remain poorly understood.

Here, we systematically charted the functional landscape of a depolymerizing TSP by constructing a comprehensive K1F DMS library and challenging it across diverse E. coli K1 hosts, directly mapping the rules of TSP function in its native phage context. This high-throughput approach yielded insights not accessible through structural or biochemical studies alone. Because this dataset spans the entire TSP, not just known substrate-binding sites, we identified functional sites outside canonical binding domains. Notably, we discovered mutations at positions such as N565 and Q615 that enhanced phage fitness despite lying outside binding sites, likely though interactions with other bacterial surface moieties. These findings highlight a broader landscape of TSP-host interactions than previously appreciated.

The dataset also offers a refined view of how the K1F TSP interacts with its polysialic acid substrate. We identified residues deep within the β-propeller active site, such as K585, that exhibited limited mutational tolerance, suggesting accommodation of longer sialic acid chains than structural assays have captured. Moreover, our results overturn some prior biochemical assignments: K410, proposed to act as a gateway residue, tolerated broad substitutions during phage infection, suggesting that another residue fulfills this role in vivo^32, 43, 46^. Beyond the active site, the β-helix stalk emerged as a “tuning knob” for processivity, where it appears to stabilize the sialic acid chain during translocation and to modulate its rate.

Comparative selections across multiple K1 hosts further demonstrated that the presence of the K1 capsule alone does not dictate infection. Instead, host discrimination arises from capsule modifications or accessory surface sugars. The β-barrel binding site emerged as a key discrimination hotspot, where certain hosts did not tolerate bulky or charged mutations to the flexible loops of the β-barrel (e.g. G730D or S460R), likely reflecting subtle differences in CPS structure or composition. The only known K1 capsule modification O-acetylation, carried out by the prophage-encoded O-acetylase NeuO, did not explain these patterns, as only one strain carried the gene, and the associated mutational profile was not observed there^57, 58, 66^.

The β-helix stalk domain was particularly important for discrimination on strains lacking O-antigen, such as EV36 which carries the rfb-50 mutation in *wbbl*^67, 68^ (Supp. 15A). Variants that destabilized sialic acid interactions were enriched on this strain, consistent with a model where faster cleavage is advantageous in the absence of O-antigen, while tighter binding is needed to overcome steric hindrance when O-antigen is present. These trade-offs between binding strength and processivity parallel what have been observed in other carbohydrate-active enzymes^69–71^. In contrast, LPS structure did not appear to contribute directly to discrimination. Although LPS has been reported as a secondary receptor for another K1 phage, the strains tested, except for one, share highly similar LPS structure, and no discriminatory patterns mapped to this feature^72^ (Supp. 15B). Together, these findings suggest that capsule and O-antigen presence, rather than LPS, are the dominant modulators of host range for K1F.

In summary, deep mutational scanning enabled the first direct functional map of a depolymerizing phage TSP in its native context. This unbiased approach uncovered both structural fragility and hidden adaptive potential, identified residues critical for host discrimination even among strains with identical capsule serotypes, and reframed the molecular basis of specificity beyond canonical binding sites. More broadly, our work establishes DMS as a powerful platform for decoding and engineering carbohydrate-active phage enzymes, paving the way for rational design of therapeutic phages with enhanced specificity and potency against encapsulated and biofilm-forming pathogens.

## Methods

### Microbes and General Culture Conditions

Bacteriophage K1F and *E. coli* EV36 were kindly provided by Dr. Antonia Sagonia. *E. coli* 10G is a lab stock originally obtained from Lucigen (60107-1). *E. coli* O45:H10 was obtained from ATCC (ATCC BAA-2649). *E. coli* A204 (AR Bank #0013, SAMN04014854) and *E. coli* A208 (AR Bank #0017, SAMN04014858) were obtained from BEI as part of the CDC *Enterobacterales* Carbapenem Breakpoint (BIT) panel.

All bacterial hosts were propagated in and plated on LB media (1% NaCl, 1% tryptophan, and 0.5% yeast extract, agar plates contain 1.5% agar and LB top agar contains 0.5% agar). Carbenicillin (100µg/mL final concentration, marker for K1F_Rec and K1F_Rec_Lib), chloramphenicol (100 µg/mL final concentration, marker for K1F_Cas9), and kanamycin (50 µg/mL final concentration, marker for K1F_Helper and T3_Helper) were added when necessary. All bacterial cultures were incubated at 37°C with shaking at 250rpm.

Microbes were stored at-80°C in 25% glycerol for long-term storage.

Bacteriophage K1F was propagated *E. coli* EV36 in LB. All phage experiments were performed using the culture conditions described above. Following propagation, phages were stored in LB at 4°C.

### Bacteria and phage propagation methods

Stationary phase cultures were created by growing bacteria for 17-20 hours overnight. Exponential phase cultures were made by diluting a stationary phase culture 1:20 and incubating until an OD_600_ of 0.6 is reached, typically taking 40min to 1 hour depending on the strain and antibiotics used. These cultures were used directly.

To determine bacterial concentrations, 100µl serial dilutions of the bacterial culture (1:10 made to 1ml up to 10^-8^) were spread on LB plates in replicate and incubated overnight. Plates with countable colonies were counted the next morning. This was used to determine concentration of each bacteria strain at an OD_600_ of 0.6 or in stationary phase and to determine CFU/mL post library transformations.

Following phage propagation on the host of interest, the lysate was purified by centrifuging at 16 *g,* and then filtering the supernatant through a 0.22µM filter. To determine titer, phage samples were serially diluted 1:10 into a final volume of 1mL LB to a 10^-8^ dilution for whole plate EOP assays. 10µl of each dilution were mixed with 250ul of relevant bacterial host in stationary phage (unless otherwise stated) and 3.5mL of 0.5% top agar and briefly vortex before plating on LB plates warmed to 37°C. After plates solidified, plates were incubated and checked for 2-6 hours and left overnight (∼15-20 hours) to establish the titer. PFU totals of 10-300 were considered acceptable. This was completed in replicate for each phage sample.

EOP was established using a reference host, usually *E. coli* EV36 with K1F_Helper unless otherwise stated. EOP values were generated by taking the phage titer on the test host divided by the phage titer on the reference host. This value was log_10_ transformed and reported as a mean of the replicates. MOI was calculated by dividing phage titer by bacterial concentration. Limit of Detection (LOD) for the K1F acceptor phage (K1F Acc) was established based on the ability to plaque on a bacterial lawn. This phage is unable to plaque on a host lacking K1F_Helper.

### Cloning Methods

All PCR reactions were performed using KAPA HiFi (<u>Roche KK2101</u>) kits. Golden Gate Assembly was performed using the New England Biosciences (NEB) Golden Gate Assembly Kit (<u>BsaI-HFv2, E1601L</u>). Phage genome assemblies were initially performed using Yeast Artificial Chromosomes (YACs) and later assemblies (clonal validations) were constructed using the NEBuilder HiFi DNA Assembly Master Mix (<u>NEB E2621L</u>). For YAC phage genome assemblies, the YAC extraction was performed using the YeaStar Genomic DNA Extraction kits (<u>Zymo D2002</u>). The Gibson Assembly Protocol (<u>NEB</u> <u>E5510</u>) using a custom lab made Gibson master mix (final concentration for the final reagents 100 mM Tris-HCL pH7.5, 20 mM MgCl_2_, 0.2 mM dATP, 0.2 mM dCTP, 0.2 mM dGTP, 10mM dTT, 1mM NAD^+^, 5% PEG-8000, 4 U/mL Taq DNA ligase, 25 U/mL Phusion polymerase, and 4U/ml T5 exonuclease) was used for plasmid based Gibson assemblies.

PCR reactions off the phage genome were performed using the typical PCR conditions however with the 95°C initial denaturation extended from 3 min to 5min. PCR reactions were performed with 1µl of undiluted phage stocks and 1µl ∼1-10ng/µl of plasmid. Following PCR reactions that used plasmid as templates, Dpn1 digestion was performed. Dpn1 digestion was carried out by adding 2% of the reaction volume of Dpn1 (<u>NEB</u> <u>R0176S</u>) and 10% reaction volume of NEB 10x CutSmart Buffer to the PCR product and incubating at 37°C for 2 hours and then heat inactivating at 80°C for 20min.

*E. coli* 10G and EV36 were transformed with plasmids and YACs using electroporation. A Bio-rad Micropulser was used for electroporation with 50µl competent cells and 1-2µl DNA (unless otherwise noted). Electroporated cells were recovered with 950µl SOC (2% tryptone, 0.5% yeast extract, 0.2% 5M NaCl, 0.25% 1M KCL, 1% 1M MgSO_4_, 1% 1M MgCL_2_, and 2% 1M glucose in dH_2_O) and then incubated at 37°C for 1 hour and plated on relevant media.

*E. coli* 10G and EV36 electrocompetent cells were made by adding 2mL overnight *E. coli* 10G or EV36 to 98mL LB (with antibiotics if necessary) and incubating at 37°C and 250rmp until an ∼OD_600_ of 0.6. Cells were then centrifuged in 5mL aliquots at 5000g for 2 minutes. The supernatant was discarded, and the cells were resuspended in 1.5mL 10% glycerol, spun down for 1 min at 5000g, and the supernatant was again discarded. This wash was repeated with 1mL and then 0.5mL 10% glycerol. Following the final wash, cells were resuspended in a final volume of 25µl 10% glycerol per tube. Method note: 5mL of culture results in 25µl of highly concentrated electrocompetent cells.

### Plasmids and engineered bacteria strains used

K1F_Helper contains a pBR backbone, kanamycin resistance cassette, mCherry, TetR, and the K1F endosialidase (*gp17*). TetR, *gp1*7, and mCherry are under constitutive expression. *Gp17* was inserted into this plasmid using Gibson assembly. K1F_Helper was used during optimized recombination and accumulation in ORACLE to prevent library bias^40^.^40^. T3_Helper contains a pBR backbone, kanamycin resistance cassette, mCherry, TetR, and the phage T3 tail fiber (also *gp17*). The T3 tail fiber was inserted into this plasmid using Gibson assembly. T3_Helper was used to test ORACLE for K1F using a non-native host system, *E. coli* 10G, and donated the T3 tail fiber in trans to the K1F acceptor phage. The K1F_Rec plasmid contains a pCDFBB backbone, an ampicillin resistance cassette, the K1F endosialidase, *gp17,* flanked by Cre lox66 sites with an m2 spacer, a 3’ pad regions, lox71 sites with a WT spacer, and a 20N barcode. The plasmid was assembled using Gibson assembly and overlap extension PCR (OEP). During assembly, OEP and site directed mutagenesis were used to remove a BsaI restriction site in WT *gp17* by introducing a synonymous mutation. This was required for subsequent Golden Gate Assembly. K1F_Rec was later used as the backbone for the DMS variant library (K1F_Rec_Lib) and used in recombination assays. Method note, while the design of the K1F_Rec plasmid was similar to that of pHRec1 used in the ORACLE method paper, a construct with both Cre recombinase and the endosialidase was not tolerated^40^.^40^. The final plasmid construct, K1F_Cas9, contains a SC101 backbone, a chloramphenicol resistance cassette, and a Cas9 cassette capable of gRNA insertion by BsaI. K1F_Cas9 with 5 gRNA derivatives was constructed using plasmid backbones with the gRNA’s already inserted, kindly provided by Phil Huss, by Gibson Assembly to swap out the spectinomycin cassette for the chloramphenicol cassette. This swap was necessary as the original plasmid construct was toxic to *E. coli* EV36. Only gRNA #3 (K1F_Cas9-3) was used during the Accumulation step of ORACLE as it was the most inhibitory to K1F Acceptor phage, all other gRNAs were used as negative controls.

EV36*^cre+^* contains Cre recombinase under tetracycline inducible expression. Cre recombinase was inserted into *fli*K of EV36 using lambda red recombination. The donor DNA containing Cre recombinase, the tetracycline inducible promoter, and the chloramphenicol resistance cassette was constructed using overlap extension PCR. Other insert sites (*RHS* and *flu*) and inducible systems (cumate and IPTG) were also tested however the *fli*K and tetracycline inducible system resulted in the best performing construct. 10G*^cre+^* similarly contains Cre recombinase under tetracycline inducible expression inserted at *fli*K using lambda red recombination. This strain was used to test ORACLE for K1F using a non-native host syste.

Plasmid backbones pBR and SC101 are lab stocks. Plasmid backbone pCDFBB was kindly provided by Dr. Claudia Schmidt-Dannert. All gene fragments used are lab stocks. All plasmids and engineered bacteria strains were sequenced using <u>Plasmidsaurus</u>.

### DMS Plasmid Library Design

To construct the endosialidase TSP DMS variant plasmid library (K1F_Rec_Lib), ∼230bp oligos were first designed and ordered from Agilent as a <u>SurePrint Oligonucleotide</u> <u>Library</u>. The designed oligos each contain a single nucleotide substitution at a single position in the entire K1F endosialidase. This included all non-synonymous substitutions, a single synonymous substitution, and a stop codon, overall resulting in a library of 22,365 variants. The most frequently found codon for each amino acid in the endosialidase was used to define each codon for each substitution. Oligos contained BsaI sites at each end for Golden Gate cloning. Due to the short length of the oligos, the library was split into 20 pools covering the length of the endosialidase. The oligo pools were amplified by PCR using recommended guidelines by Agilent. The K1F_Rec plasmid was used as template for PCR to create 20 backbones for each of the pools. A 20N barcode, later required for variant barcode mapping, was also constructed using a primer with 20N random nucleotide as a template for PCR. Each oligo pool, corresponding backbone, and a 20N barcode were assembled using Golden Gate Assembly with 0.075pmol backbone, 0.15pmol oligo insert and 0.15pmol 20N barcode insert. This reaction mix was then cycled from 37°C to 16°C over 5 minutes, 30x, then held at 60°C for 5 minutes to complete the Golden Gate Assembly. Membrane drop dialysis was then performed for 1 hour to enhance transformation efficiency. Next, the entire reaction volume (∼20µl) was transformed into 100µl *E. coli* 10G cells. The transformed cells were than plated at dilutions ranging from 10^-1^ to 10^-3^ to determine the total number of cells. 1mL of each pool or a 10^-1^ dilution were added to 100mL LB and grown for 17 hours overnight. The overnight resulting in ∼2 x 10^5^ CFU/mL transformed cells, as determined by the dilution plating, had the plasmids purified. This number of transformed cells was targeted as it would result in ∼150 barcodes per variant which is required for subsequent variant barcode mapping. Plasmid concentrations for each of the 20 pools were determined by nanodrop, and then each pool was combined at an equimolar ratio to create the final plasmid variant pool (K1F_Rec_Lib). K1F_Rec_Lib was then transformed into 14 100ul aliquots EV36*^cre+^* + K1F_Helper, recovered in a final volume of 1mL for one hour for each transformation. Following recovery, dilutions, ranging from 10^-1^ to 10^-3^, of all 14 transformations were pooled together at a 1:1 ratio and initially plated on LB plates containing carbenicillin, chloramphenicol, and kanamycin to estimate transformation efficiency. The total of actual transformed cells was estimated to be 1 x 10^5^ CFU/mL. The remainder of the 14ml pooled library was added to 1.4L of LB with chloramphenicol, kanamycin, and carbenicillin and incubated for 17 hours overnight. 14 separate transformations were used for full library transformation due to the lower transformation efficiency of EV36*^cre+^ +* K1F_Helper and the number of transformants as determined by the number of transformed cells of one 100µl transformation reaction. This host, EV36*^cre+^* with K1F_Helper and K1F_Rec_Lib was the host used for Optimized Recombination during ORACLE.

### Plasmid Library Barcode Mapping

Variant barcode mapping was required for higher read depth and use of high fidelity short read Illumina NGS instead of long read sequencing methods to measure library member abundance across the endosialidase. Barcode mapping for all 20 library pools was accomplished by creating separate barcode-mapping constructs for each pool. The separate barcode-mapping constructs were assembled using a one-part Golden Gate Assembly to place the 20N barcode directly adjacent to library pool of interest. This was then followed by NGS to determine barcodes associated with each variant. For the Golden Gate Assembly, K1FRec_Lib, throttled to have ∼150 barcodes per variant as described above, was PCR amplified with primers that introduced Bsa1 sites at the end of the target oligo pools and in front of the associated 20N barcode. This was then assembled using the NEBridge BsaI Golden Gate Assembly kit, 0.075pmol backbone, and 2µl T4 DNA Ligase buffer. This reaction mix was then cycled from 37°C to 16°C over 1 minute, 30x, then held at 60°C for 5 minutes to complete the Golden Gate Assembly. Membrane drop dialysis, using a 0.22µm membrane, was then performed for 1 hour to enhance transformation efficiency. Next, the entire reaction volume (∼20ul) was transformed into 100µl *E. coli* 10G cells and recovered at a final volume of 1mL for 1 hour. The transformed cells were than plated at dilutions ranging from 10^-1^ to 10^-3^ to determine the total number of transformants. The remainder of the transformation reaction was then added to 100mL LB + carbenicillin and incubated for 17 hours overnight. The total of actual transformed cells was estimated to be in the range of 1-5 x 10^5^ CFU/mL across all 20 pools. All libraries were sequenced using Illumina sequencing platforms.

### Recombination Rate and Accumulation Assays

To determine the recombination rate, K1F acceptor phage was passaged on 5mL exponential phase *E. coli* EV36*^cre+^* with K1F_Helper and K1F_Rec that had been induced for 1.5 hours with 0.5uM tetracycline. K1F_Rec was used because it carries the WT endosialidase which ensures that every recombined phage is able to plaque with equal efficiency. An MOI of 10 and passage of 30 minutes was used for the assays so to evaluate the recombination rate after only one passage on the host. Phages were then purified to determine the final phage population. Recombination rate was determined by dividing the phage titer following plating on EV36 containing K1F_Helper by that seen when plated on EV36. Only recombined phages are able to plaque on EV36 as the K1F acceptor phage does not have the endosialidase in there genome.

The accumulation of recombined phages over acceptor phages in the Accumulation step of ORACLE was validating using the exact protocol as described previously^40^.

### ORACLE Steps

All ORACLE steps were completed similar to what was described by Huss et al. with a few key differences that will be highlighted here^40^. The first iteration of the K1F Acceptor phage was constructed using YAC rebooting as described previously, however, all other phage constructs were assembled using the NEBuilder HiFi DNA Assembly Master Mix^65^.

The most distinct difference from the previously described ORACLE steps can be observed in the Optimized Recombination step as Cre recombinase is under inducible control in the bacterial host genome. For the Optimized Recombination step of ORACLE, a 1:40 back dilution of an overnight culture of EV36*^cre+^* with K1F_Rec_Lib and K1F_Helper with a final volume of 25mL was made. This back dilution was allowed to recover for 30 minutes before adding 0.5µM tetracycline to induce Cre recombinase production. Following induction, the culture was grown to an OD_600_ of 0.6 (∼1.5 hours) before the addition of K1F Acceptor phage at an MOI of ∼10. The cultures were incubated until lysis (∼40 minutes) and purified. This library phage population constitutes the initial phage population and contained 5.98 x 10^7^ variants/mL in a total phage population of ∼9.6×10^10^ PFU/mL. The remainder of the phages in the population are acceptor phages. For the next step, Accumulation, there are also key differences between what was done here and what was described previously. This step was performed by adding an MOI of 0.1 of recombined phages to a 25mL culture of exponential phase EV36 with K1FHelper and K1F_Cas9. Cultures were incubated until lysis (∼2 hours) and then purified. Exponential phase was chosen over stationary phase, used in the original method, as cultures did not lyse during this step when stationary phase was used. Furthermore, exponential phase was more inhibitory based on EOP. The Library Expression step of ORACLE was performed as described previously^40^.

### DMS Selection

DMS selection was performed the same across all bacteria strains that readily lysed upon addition of library, including *E. coli* EV36 and all additional multi-drug resistant *E. coli* strains tested. The K1F TSP variant library was added to 5mL of exponential host at an MOI of ∼0.01 and the culture was allowed to fully lyse. The phage lysate was then filtered purified and prepared for sequencing. The MOI of 10^-2^ was chosen to allow for the greater enrichment of winners or losers in the population due to the large library size and to allow slower growing phages a chance to replicate. At this MOI, we expect wildtype to complete four total infections cycles. The entire selection and purification process was repeated in biological replicate for each host.

### Next Generation Sequencing Preparation

NGS (deep sequencing) was used to map barcodes to variants and to evaluate phage populations following selection. For variant-barcode mapping, NGS samples were prepared using 20ng of Golden Gate product that was produced as described above for each of the 20 pools. Two rounds of PCR were used to amplify the final NGS products. The first round of PCR used primers that bind directly 5’ to the library pool of interest and into a constant region 3’ of the 20N barcode. These primers add a variable N region that is necessary to assist with nucleotide diversity during sequencing and the universal Illumina adaptor. This first round of PCR was performed using 12 total cycles and the resulting product purified. The second PCR reaction adds an index and the Illumina “stem” and uses 1µl of the purified first round product as the template for the reaction. This reaction was carried out for 9 cycles and the resulting purified product directly used for sequencing. All barcode mapping samples were sequenced using an Illumina NextSeq system in our lab, 2 x 150bp read length using NextSeq Reagent Kit with a P2 flow cell and 300 cycles or sent for shared NovaSeq sequencing to the <u>University of Wisconsinbioinformatics core</u>. For evaluating phage populations following selection, a similar approach was used for PCR amplifications except that PCR was performed directly off pure phage lysate. First round primers bind directly adjacent to the 20N barcode in constant regions flanking either side and again introduce a variable N region. Following the second round of PCR, the products were purified and directly sequenced using NextSeq Reagent Kit with 2 x 50bp read length and a P2 flow cell with 100 cycles or sent for NovaSeq sequencing at the UW bioinformatics core.

### Next Generation Sequencing Analysis for Plasmid Barcode Variant Mapping

Paired-end Illumina sequencing reads were merged and quality filtered using fastp, a FASTQ preprocessor^73^. The output sequences were trimmed to have only the pool of interest and the 20N barcode. The nucleotide sequences of the variant pool were translated, and each variant called using a custom script. Only single substitution variants were kept. The final output was a list of each variant with ∼200 associated barcodes per variant, with a few exceptions. To ensure barcodes were properly mapped, a higher read cutoff (10-15 depending on read coverage) was used. Additionally, if a barcode constant was seen associated to more than one variant, it was only kept if it passed a quality filter value called a “chastity value” of greater than 0.98. This value was calculated by taking the highest abundance value for the target barcode and dividing by all other abundance values for that same barcode mapping to different variants. For example, if the abundance value for one barcode is 150 and 50 for the same barcode associated with a different variant, the chastity value would be 50/150=0.33 and the barcode would be thrown out. Following the variant barcode mapping for all 20 pools, there is no discernible skew, and 95.2% of the full library (21,289 variants of the 22,365 member library) was present with at least 1 barcode mapped. Following barcode variant mapping, an output containing a table of all variants and their associated barcodes was generated. This table was later used to map variants back to associated barcodes following K1F DMS library sequencing.

### Next Generation Sequencing Analysis for Phage Library DMS Selection

The paired-end Illumina sequencing reads were merged, and quality filtered using the same methods as described for the plasmid barcode variant mapping. The cleaned output sequences were then trimmed to only contain the 20N barcode sequence. The barcode sequences and corresponding counts were then mapped to the variant-barcode table that was generated previously (see above) to determine associated variants using a custom script. To filter for possible sequencing errors that would result in barcode counts associated with the wrong variant, we removed all barcodes mapped to multiple variants and ensured that each barcode was reporting similar fitness trends for the associated variant following library selection (see blow section). To avoid missing low abundance members, a read cutoff of 4 for all post-selection NGS analysis was used.

The pre-selection K1F DMS library contained 85% of the 22,365 library variants. The missing variants were excluded from subsequent analysis as a pre-selection population could not be accurately determined, although some missing variants emerged in post-selection populations. Biological replicates were correlated to determine reproducibility.

The pre-selection library showed skew, however, library member abundance correlated well between biological replicates indicating skew was likely introduced by lack of barcode coverage and read depth for the missing replicates rather than skew created during the DMS library creation.

### Functional Score Calculation

To score enrichment for each variant, a basic functional score (F) was used. The functional score for each variant were calculated by taking the variant abundance post selection and dividing by the variant abundance prior to selection. Variant abundance pre and post selection was calculated by summing up read counts for each barcode associated with a given variant. Prior to summing up read counts for all barcodes, outlier barcodes were removed from the dataset. To identify outlier barcodes, functional scores were calculated for each individual barcode associated to a given variant and then compared across all barcode reported functional scores for the variant. Barcodes reporting functional scores with a z-score greater than 2 were removed from the dataset. Furthermore, all barcodes mapping to more than one variant following selection were removed from the dataset. To compare functional scores across hosts, each functional score was normalized to wild-type to yield the normalized functional score (Fn).

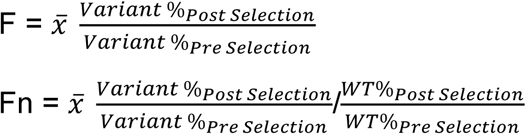

To calculate the net impact functional score for each position across the K1F TSP, log_2_ transformed functional scores of all variants for a given position were summed. Variants that dropped out following selection were given a limit of detection (LOD) functional score of-3. Net impact functional scores greater than 0.8 were labeled “positive impact”, scores less than 0.8 but greater than-0.3 were labeled “no impact”, and any net functional scores less than-0.3 were labeled “negative impact”.

### Hierarchical Clustering

Hierarchical clustering was performed in R using the libraries “cluster” and “factoextra”. The clustered heatmap was then generated in R using the libraries “pheatmap”, “ComplexHeatmap”, “gplots”, “dendextend”, and “circlize”. The correct cluster count was determined using the gap statistic, elbow method, and silhouette method using the R libraries “factoextra” and “cluster”. The Gap Statistic provided the most accurate report of the correct cluster count (see relevant supplemental figures). The correct clustering method was determined using the cophenetic correlation coefficient. The cophenetic correlation coefficient for each clustering method was calculated in R using libraries “cophenetic” and “dendextend”. The correct linkage method was determined using the agglometative coefficient which was calculated in R using the library “cluster”. Based on these metrics, the optimal distance measure was determined to be Spearman and the best linkage method Ward.

### Clonal Validations

K1F clonal variants were constructed using Gibson Assembly and the NEBuilder HiFi DNA Assembly Master Mix. OEP and site directed mutagenesis was used to introduce the desired mutation into the endosialidase for each clonal variant. Following Gibson Assembly, membrane drop dialysis was performed for ∼1 hour on the entire reaction volume. Next, 50µL *E. coli* 10G electrocompetent cells were transformed with the entire reaction volume and recovered in a total volume of 1mL SOC for 1 hour at 37°C. *E. coli* 10G was used for transformation versus the phage K1F host EV36 due to the higher transformation efficiency. Following recovery, 40µL of chloroform was added to each transformation, vortexed for 10 seconds, and incubated for another 20 seconds. The vortexing and incubations steps were repeated 3 times before spinning each culture down for 2 minutes at 10,000 rpm. The supernatant was then removed and filter purified through a 0.22µm syringe filter. This chloroform treatment was required to release the assembled phages from the cells. The resulting purified phage lysate was then subjected to whole plate EOP assay on EV36 + K1F_Helper and incubated overnight at 37°C. Individual plaques were then isolated, propagated on exponential phase EV36 + K1F_Helper and filter purified through a 0.22µm filter. The genomes were then extracted using the Norgen Phage DNA Isolation Kit (<u>Norgen #46800</u>) and sent for sequencing.

For clonal validations, each K1F clonal variant was subjected to EOP assays on target hosts using EV36 + K1F helper as the reference host. EOP assays were conducted as described above.

### Genomic Validations

The genomic sequences of *E. coli* EV36, *E. coli* A204 (AR Bank #0013), and *E. coli* A208 (AR Bank #0017), were obtained were obtained from NCBI with accession numbers CP079993.1, NZ_CP032204.1, and NZ_CP024886.1 respectively. The genomic sequence of *E. coli* O45:H10 (ATCC BAA-2649) was obtained from ATCC. Finally, the genomic sequence of UTI50 was obtained from in house sequencing on an Illumina NextSeq platform and assembled using SPAdes (https://github.com/ablab/spades). All genomes were then annotated using Prokka and Rast-tk to ensure comprehensive gene predictions. Pangenome analysis across these genomes was performed using Panaroo. This enabled the identification of the presence and absence of polysialic acid genes among these genomes.

The O-antigen gene cluster was extracted by isolating the genomic region between the gnd and galF genes. Since the EV36 genome lacks the galF gene, a 17 kbp region downstream of the gnd gene was extracted as the O-antigen gene cluster. Similarly, the lipopolysaccharide (LPS) core biosynthesis gene cluster and KPS gene cluster were identified by extracting the gene regions spanning from waaA to waaD and kpsM to kpsF, respectively. A Python script was developed to automate the extraction of these gene clusters from the annotated genomes.

## Acknowledgements

We thank R. Welch for UTI strains and A. Sagonia for bacteriophage K1F and *E. coli* EV36. We also thank Dr. Chutikaarn Chitboonthavisuk and Rebecca Back for helpful discussion on data analysis and troubleshooting, and Dr. Monica Neugebauer for insights into enzyme structural analysis. Finally, S.E. acknowledges support from the Steenbock Predoctoral Fellowship and the William H. Peterson Fellowship, awarded by the University of Wisconsin-Department of Biochemistry, which made this research possible.

## Author contributions

Conceptualization: S.E., P.H., and S.R. Methodology: S.E., P.H., and S.R., Investigation: S.E. and D.K.K.B., Visualization and Data curation: S.E, Software: S.E. and P.H., Writing (original draft): S.E., Writing (review and editing): S.E., P.H., and S.R., Resources: S.R. and K.A., Supervision: S.R., Funding acquisition: S.R.

