## Supplementary_Figures for "Mapping Structural Constraints and Adaptive Potential in a Capsule-Degrading Phage Tailspike Protein"

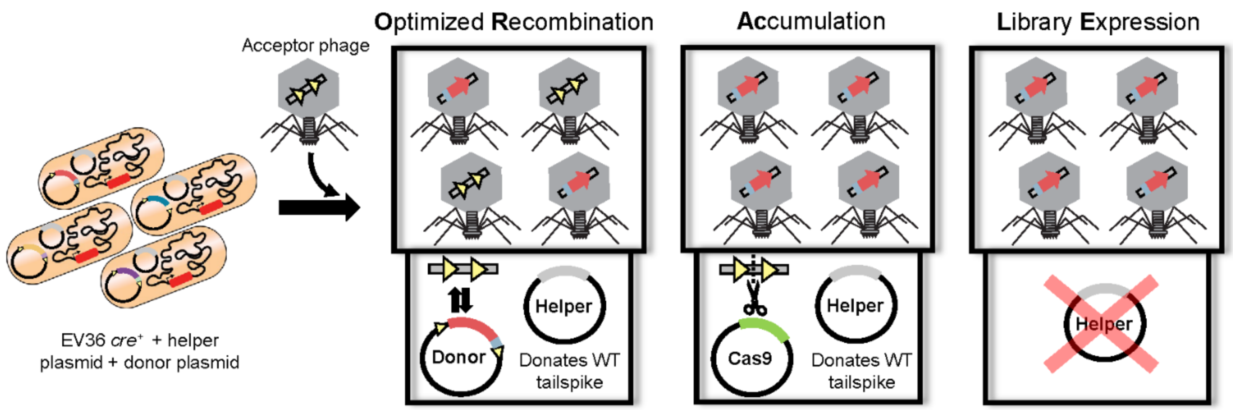


**Supplementary Figure 1**. Modified ORACLE workflow to generate the K1F tailspike protein deep mutational scanning library. EV36 *cre*$^{+}$ is *E. coli* EV36 carrying Cre recombinase integrated into the genome. Scissors indicate the targeted cleavage of the acceptor sequence by Cas9.

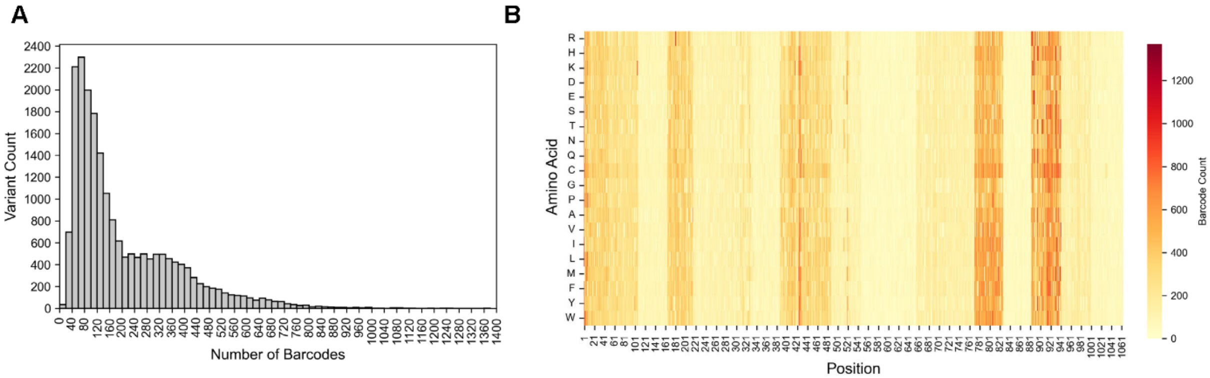


**Supplementary Figure 2.** Distribution of plasmid-mapped barcodes. (**A**) Histogram of the number of barcodes per variant in the plasmid library pool prior to insertion into the phage genome. (**B**) Heatmap of the barcode count for each variant at each position. Stripes across the heatmap indicate cutoff for individual assembled library pools.

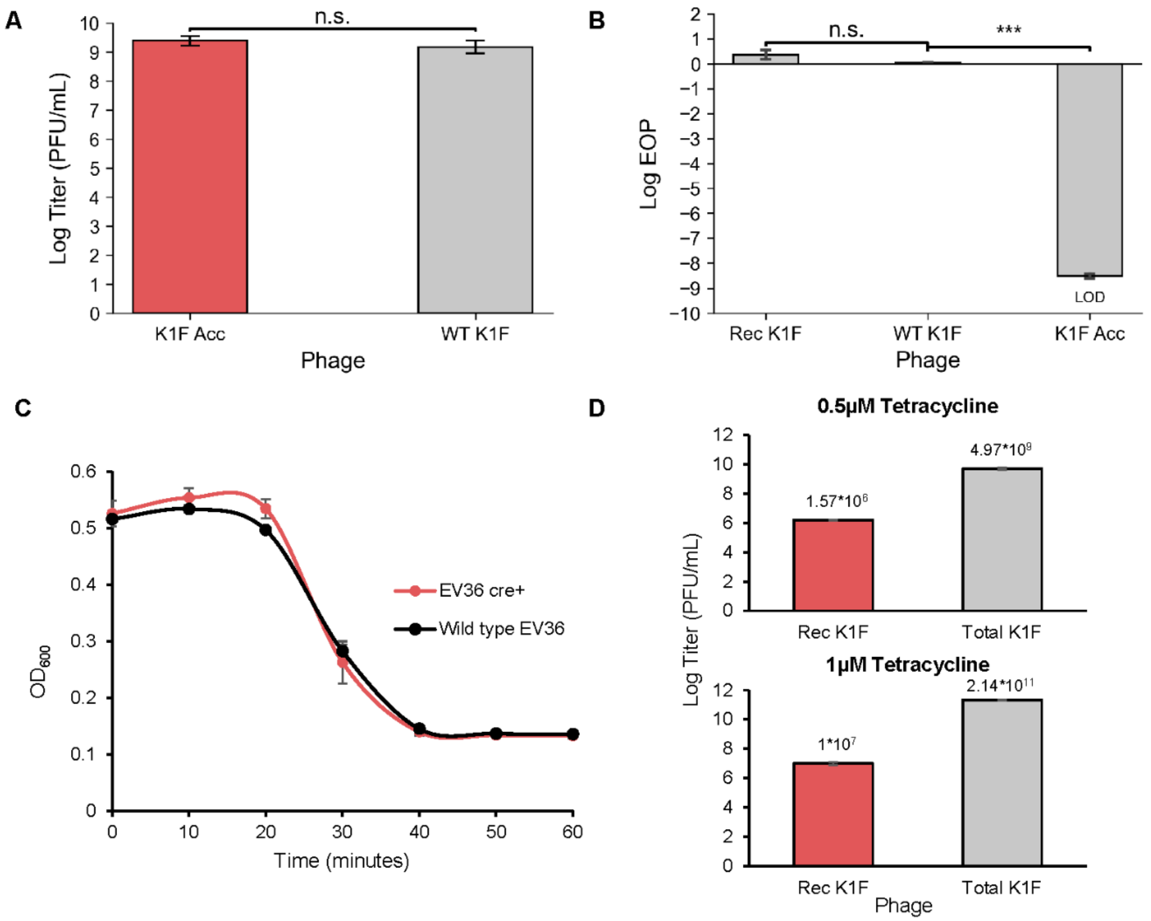


**Supplementary Figure 3**. Each step of ORACLE can be adapted to phage K1F. (**A**) Plaque assay results for wild type K1F (WT K1F) and K1F acceptor (K1F Acc) on EV36 carrying the K1F_Helper plasmid. No plaquing deficiency is observed ($p=0.26$). n.s. stand for not significant. (**B**) Efficiency of plating (EOP) assay on wild type EV36 for recombined K1F with the wild type tailspike protein inserted into the genome (Rec K1F), wild type K1F (WT K1F), and K1F acceptor phage (K1F Acc). n.s. $p > 0.05$, *** $p < 0.001$. LOD is limit of detection. (**C**) Growth curve of EV36 with Cre recombinase integrated into the genome (EV36 cre$^{+}$) and wild type EV36 upon infection by wild type K1F. (**D**) Titers of recombined K1F phage (Rec K1F) versus Total K1F phage following the Optimized Recombination step of ORACLE using two different induction conditions. Recombination efficiency for 0.5μM was $3.16*10^{-4}$ and for 1μM was $4.67*10^{-5}$.

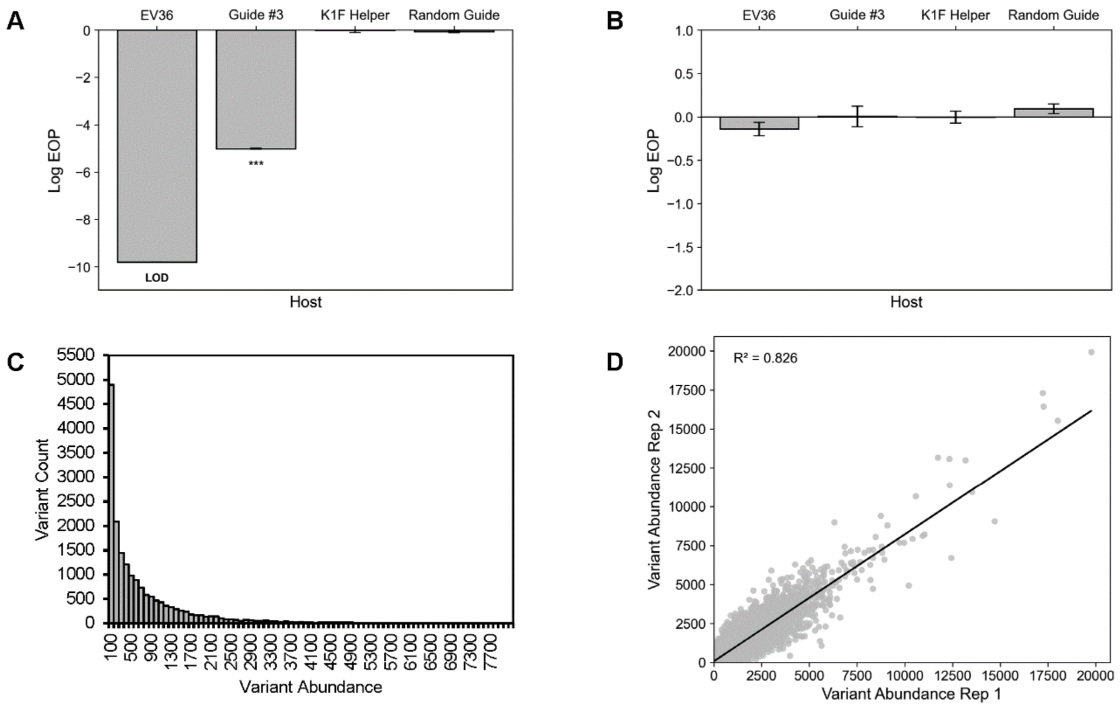


**Supplementary Figure 4**. Each step of ORACLE can be adapted to phage K1F. (**A**) EOP assay K1F acceptor phage across wild type EV36, Guide #3, and Random Guide. EV36 + Helper is the reference stain. LOD stands for limit of detection, *** $p < 0.001$. (**B**) EOP assay for recombined K1F phage across wild type EV36, EV36 carrying the K1F_Helper plasmid (EV36 + Helper), EV36 carrying the sgRNA guide #3 Cas9 plasmid and K1F_Helper plasmid (Guide #3), and EV36 carrying a randomized sgRNA Cas9 plasmid and K1F_Helper plasmid (Random Guide). EV36 + Helper is the reference stain. (**C**) Histogram of library member abundance following the library expression step of ORACLE. (**D**) Scatter plot showing correlation between variant abundance of two replicates of the K1F tailspike protein library following the ORACLE library assembly process.

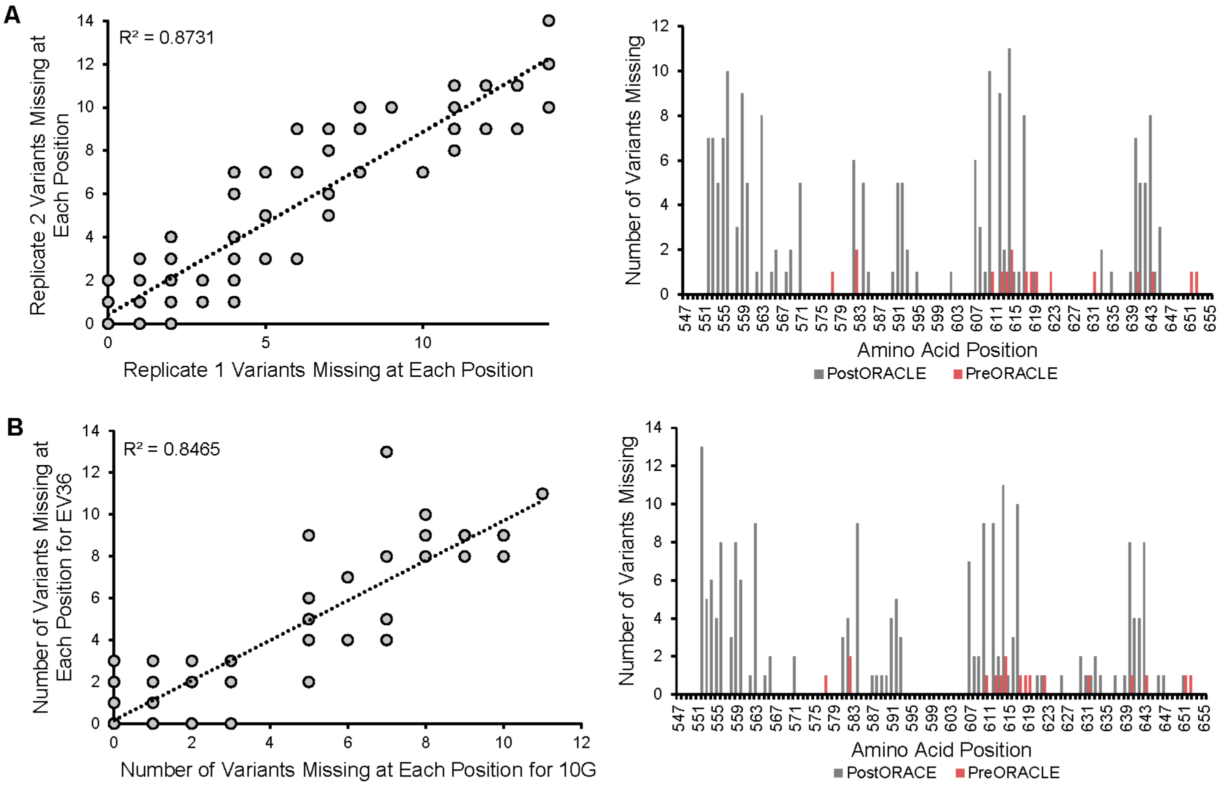


**Supplementary Figure 5**. Library bias is caused by negative selection during library assembly. (**A**) Left, a scatterplot of the correlation of number of variants missing at each position between replicates following ORACLE for a 2,268-member K1F library. Right, another representation of the number of variants missing at each positions pre- and post- ORACLE, (**B**) Left, a scatterplot of the correlation of number of variants missing at each position following ORACLE using EV36 versus non-native host *E. coli* 10G for a 2,268-member K1F library. Right, number of variants missing at each position pre- and

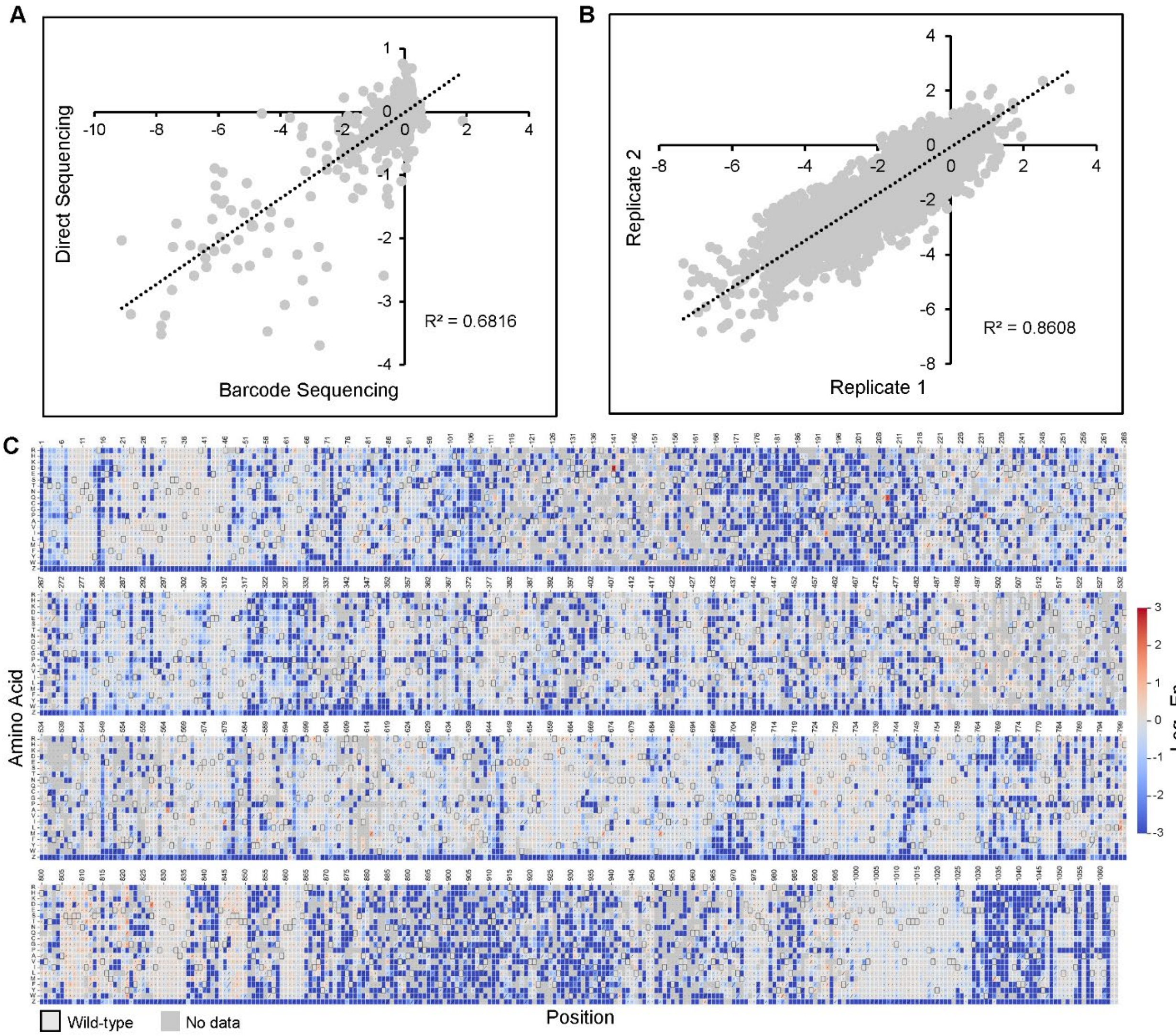


**Supplementary Figure 6**. Normalized functional scores (Fn) reported following library selection on *E. coli* EV36. (**A**) Correlation of Fn scores between direct sequencing of a ~250 bp region of the K1F tailspike protein (TSP) versus barcode sequencing. (**B**) Correlation of Fn scores between K1F TSP library biological replicates following selection on *E. coli* EV36. (**C**) Heatmap of the Fn scores of each variant following selection on *E. coli* EV36. Limit of Detection (LOD) is Fn= -3. Diagonal lines in squares represent standard deviation.

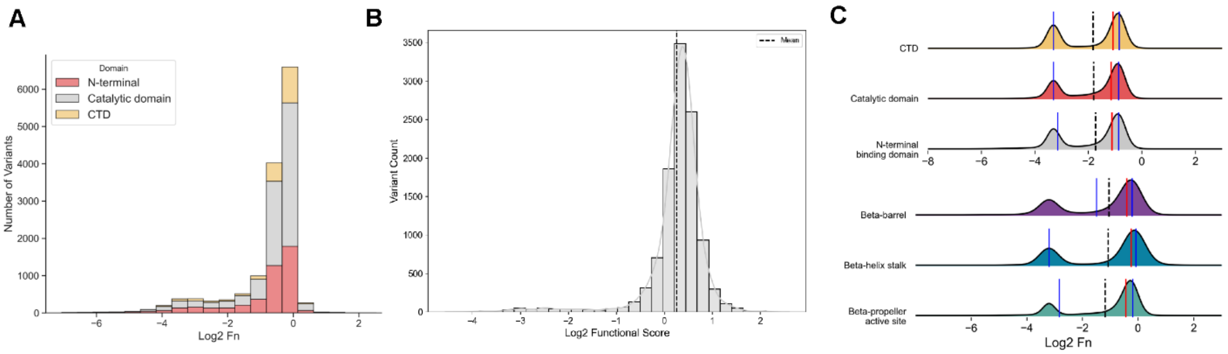


**Supplementary Figure 7**. Functional distribution across different domains/sites of the K1F tailspike protein following selection on *E. coli* EV36. (**A**) Histogram of the normalized function score (Fn) distribution colored by domain. Variants that dropped out following selection are not included. (**B**) Functional score distribution of wild type barcodes. Standard deviation is 0.6 and the mean in 0.2. (**C**) Ridgeline plot for the Fn across domains/sites of the K1F TSP. The dashed line indicates the mean score. Red line is the median, and blue lines interquartile range (25th and 75th percentiles).

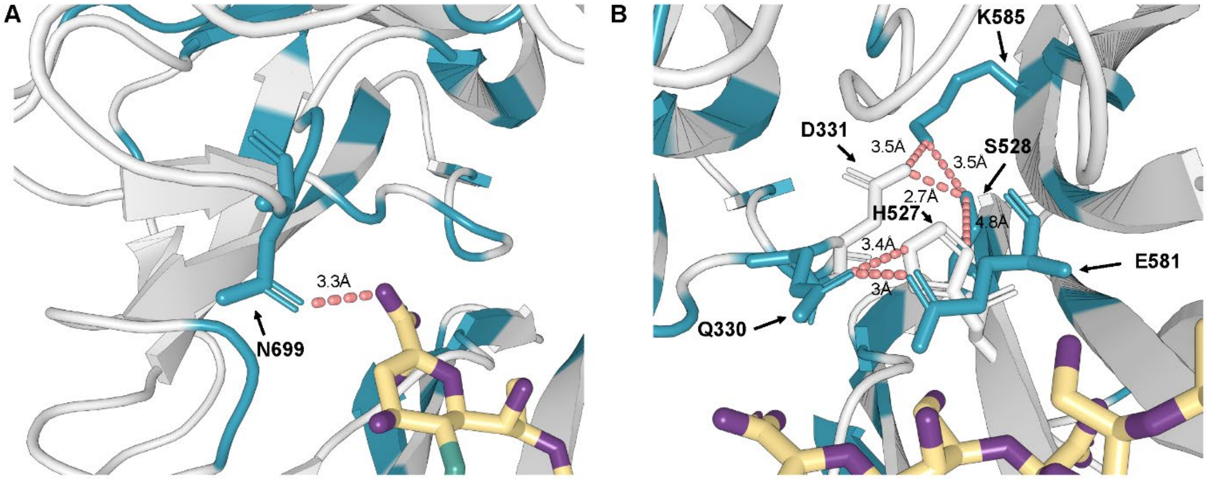


**Supplementary Figure 8**. Novel interactions with substate and catalytic active site residues following K1F tailspike protein (TSP) library selection on *E. coli* EV36. (PDB: 3gvk) (**A**) PyMol structure of position N699 forming hydrogen bonds with the sialic acid chain in the β-propeller active site. The sialic acid chain is colored gold in licorice representation. Oxygens are colored purple and nitrogen colored green. The hydrogen bond is represented by red dashes. (**B**) Polar channel formed by position K585 interaction with E581 in the β-propeller active site.

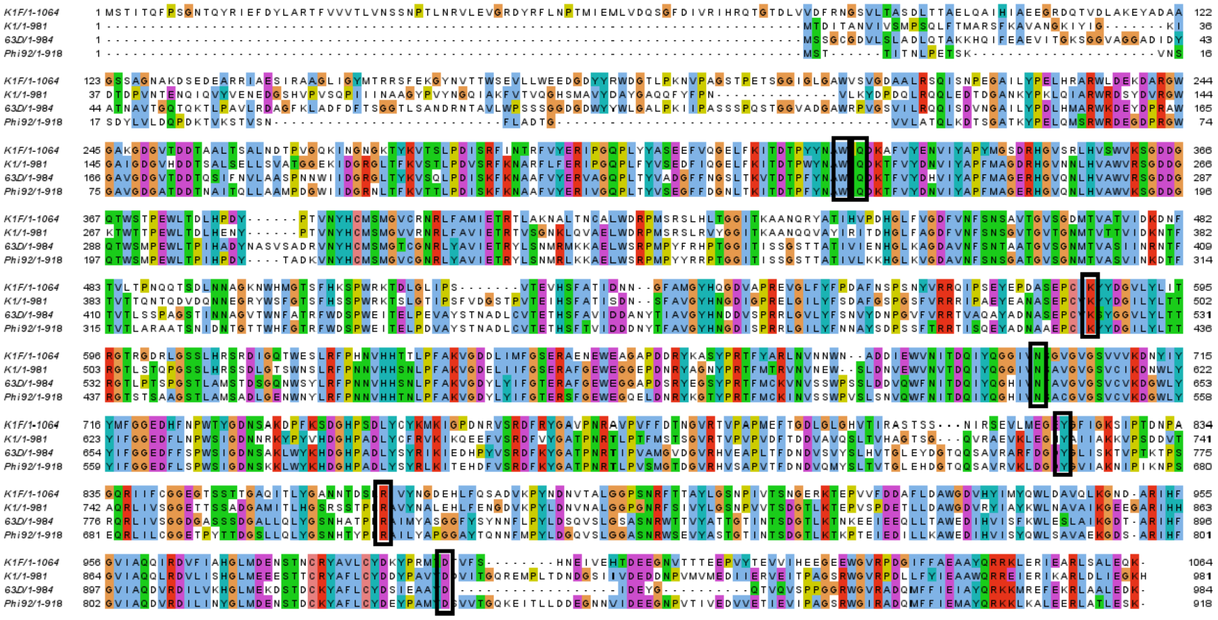

**Supplementary Figure 9**. Sequence alignment using Clustal to the K1F endosialidase of four previously characterized endosialidases from phages; K1, 63D, and Phi92. Conserved positions are colored. Newly discovered and conserved positions are boxed in black (W328, Q330, K585, N699, Y821, R866, and D994).

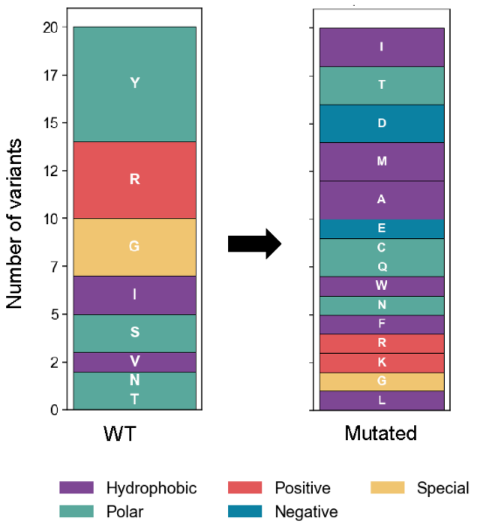


**Supplementary Figure 10**. Distribution of mutation types for the top 20 performing variants in the β-helix stalk. Mutations are colored by properties and ordered by abundance.

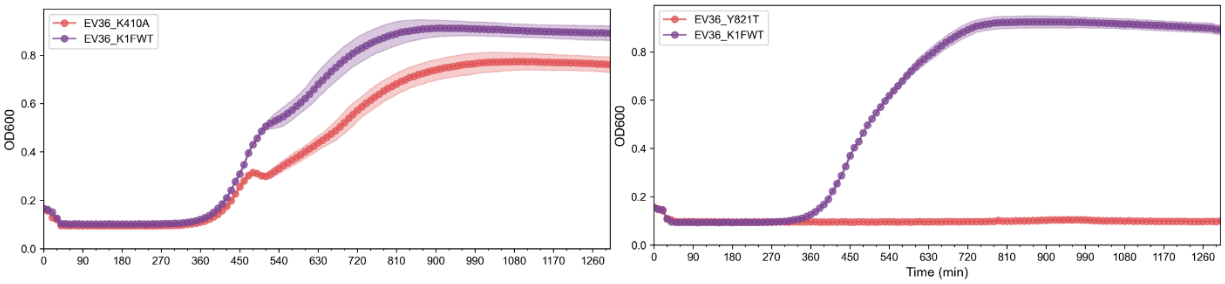


**Supplementary Figure 11**. Growth curves for the two clonal validation variants, K410A and Y821T, passaged on EV36 compared to wild type K1F (K1FWT).

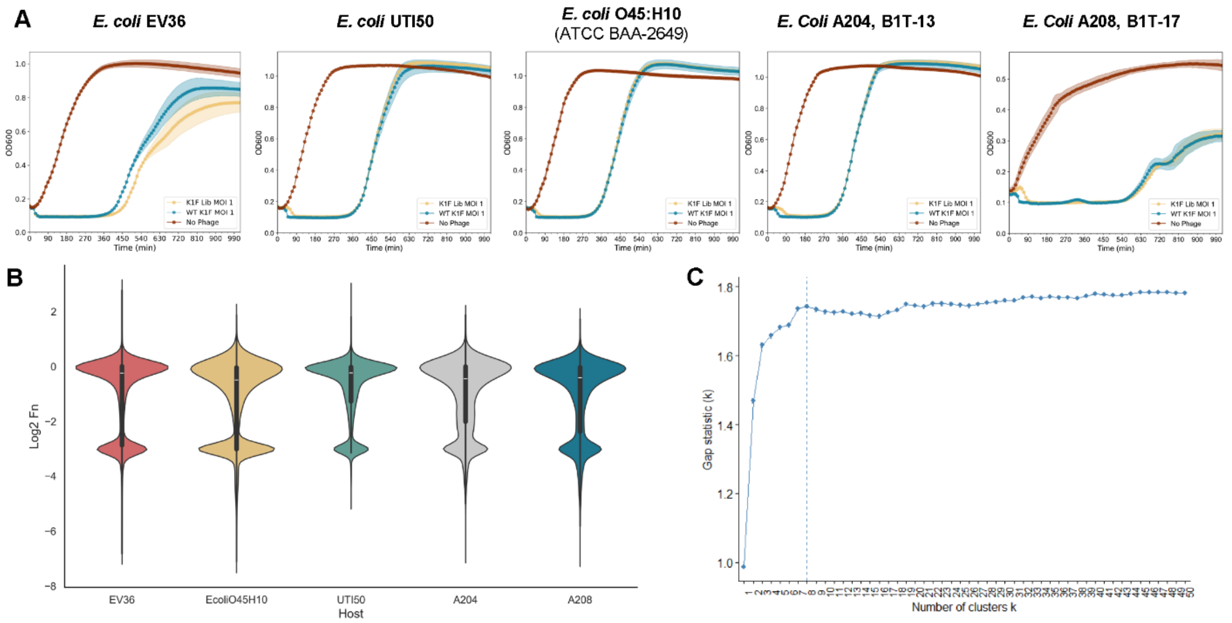


**Supplementary Figure 12**. Selection of the K1F tailspike protein (TSP) DMS library on a diverse set of *E. coli* K1 strains yields different functional profiles. (**A**) Growth curves for each strain with no phage (brown), wild type K1F (WT K1F) at an MOI of 1 (blue), and the K1F TSP library (K1F Lib) at an MOI of 1 (yellow). (**B**) Violin plots of the Log2 normalized functional scores following selection of the K1F TSP library on each strain. (**C**) Scatter plot of the Gap statistic (k) for different numbers of clusters.

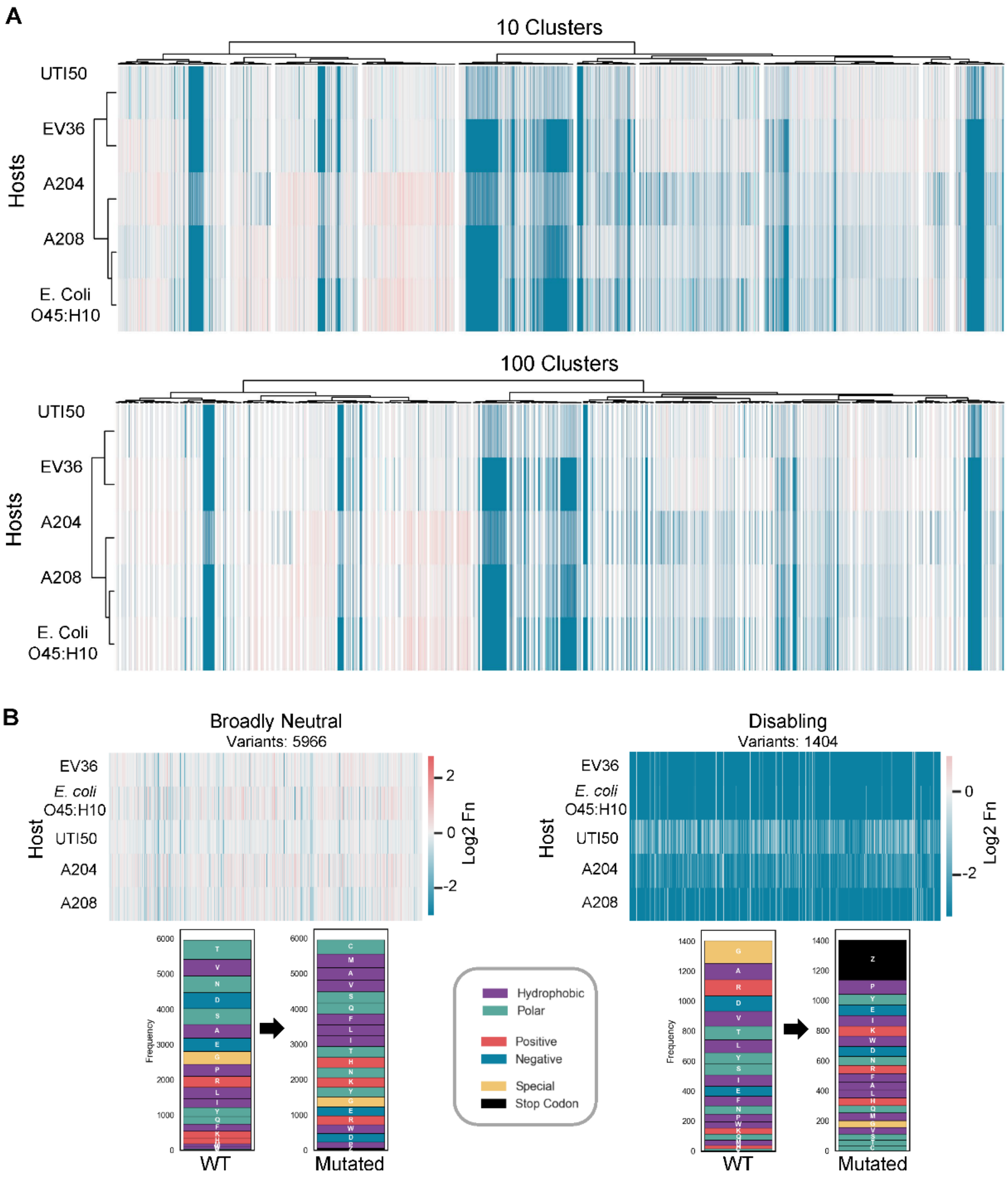


**Supplementary Figure 13**. Different cluster cutoffs do not change the relationships between each variant and host (**A**) Hierarchical clustering heatmap for functional scores following selection of the K1F DMS library on each of the 5 bacterial strains with 10 clusters and 100 clusters. (**B**) Heatmaps for all broadly neutral and disabling clusters. Below each heatmap, there is a representation for the total abundance of all wild type amino acids and what they were mutated to stacked from most abundant to least abundance. Each amino acid is colored by property.

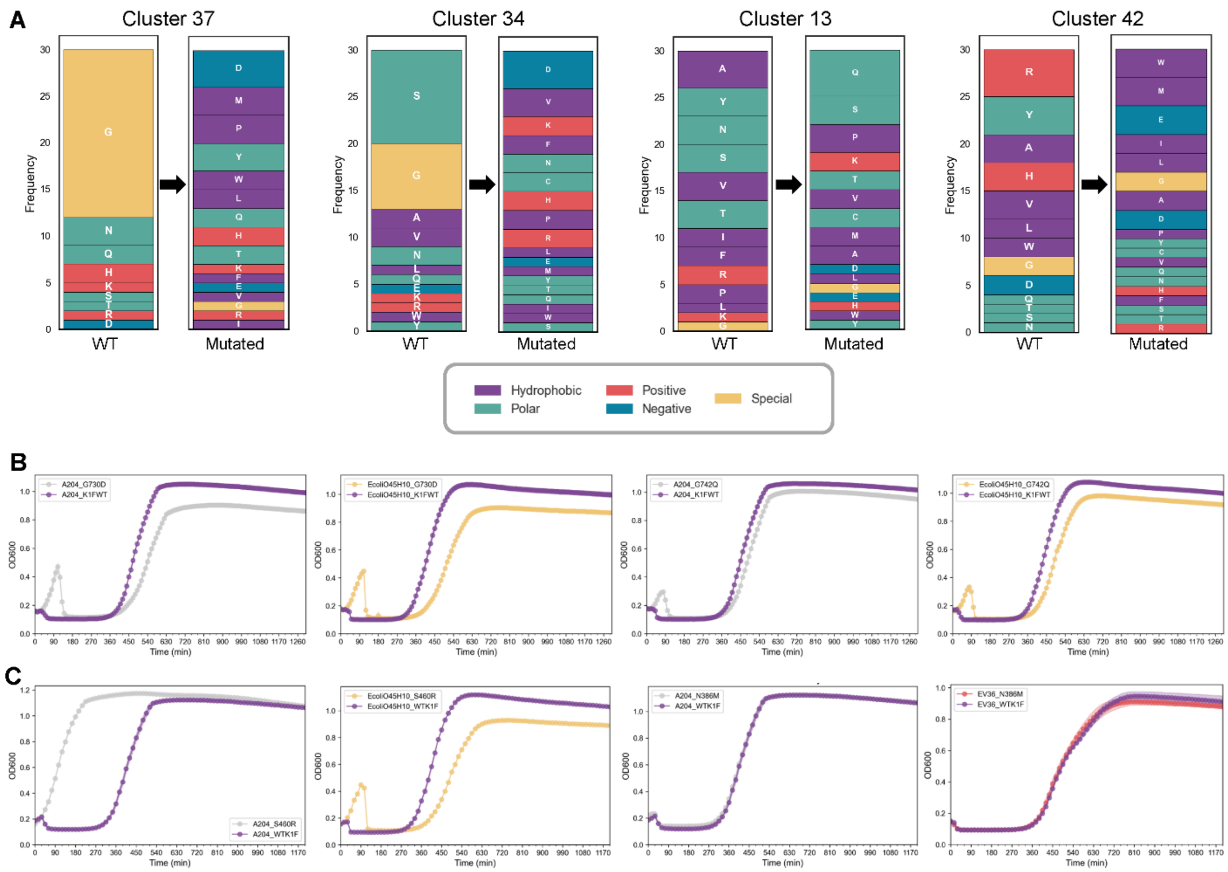


**Supplementary Figure 14**. Novel host discriminating positions can be discovered upon hierarchical clustering. (**A**) Top 30 discriminating variants on 2 hosts (Clusters 37 and 34) or on 1 host (Clusters 13 and 42) compared to all other hosts. The wild type amino acids are on the left and the mutations are on the right. The amino acids stacked by abundance and colored by property. (**B**) Growth curves for the two clonal variants, G730D and G742Q, passaged on the two discriminatory hosts A204 and *E.coli* O45:H10 compared to wild type K1F (K1FWT) in replicate. (**C**) Growth curves for the two clonal variants S460R and N386M passaged on the two discriminatory hosts A204 and *E.coli* O45:H10 . A growth curve for N386M passaged on EV36 is shown as a reference.

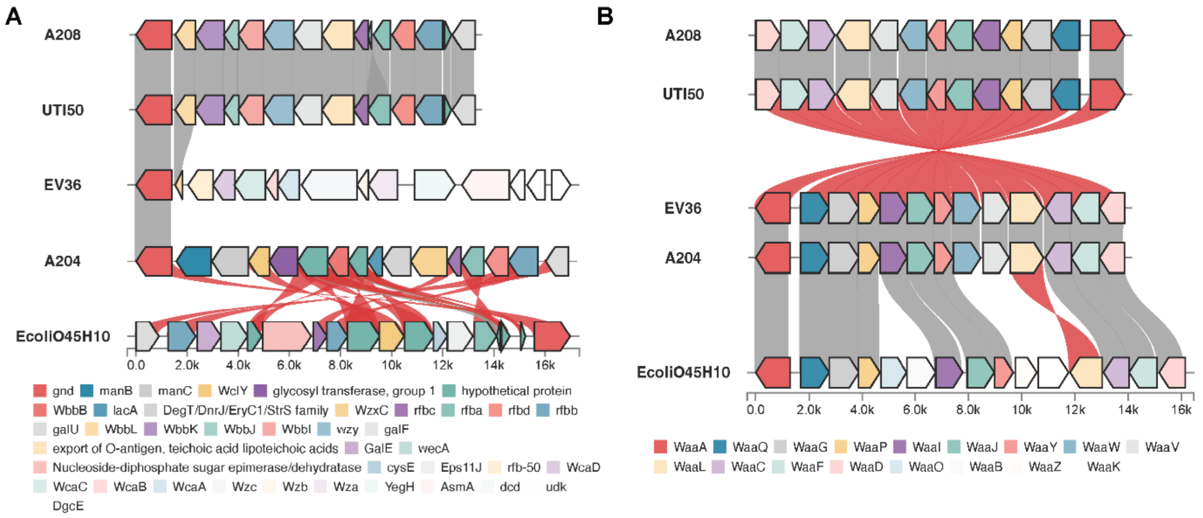


**Supplementary Figure 15**. Genomic validations reveal that differences in the bacterial surface sugars don't necessarily explain observed host discrimination. (**A**) Comparison of the O-antigen gene cluster across the five screened bacterial strains. Shaded lines connecting genes indicate shared genes across strains. (**B**) Comparison of the lipopolysaccharide gene cluster across the five screened bacterial strains. Shaded lines connecting genes indicate shared genes across strains.
